# The power of geohistorical boundaries for modeling the genetic background of human populations: the case of the rural Catalan Pyrenees

**DOI:** 10.1101/2022.10.28.513229

**Authors:** Joan Fibla, Iago Maceda, Marina Laplana, Montserrat Guerrero, Miguel Martín Álvarez, Jesús Burgueño, Agustí Camps, Jordi Fàbrega, Josefina Felisart, Joan Grané, José Luis Remón, Jordi Serra, Pedro Moral, Oscar Lao

## Abstract

The genetic variation of the European population at a macro-geographic scale follows genetic gradients which reflect main migration events. However, less is known about factors affecting mating choices at a micro-geographic scale. In this study we have analyzed 726,718 autosomal SNPs in 435 individuals from the Catalan Pyrenees covering around 200 km of a vast and abrupt region in the north of the Iberian Peninsula, for which we have information about the geographic origin of all grand-parents and parents. At a macro-geographic scale, our analyses recapitulate the genetic gradient observed in Spain. However, we also identified the presence of micro-population substructure among the sampled individuals. Such micro-population substructure does not correlate with geographic barriers such as the expected by the orography of the considered region, but by the bishoprics present in the covered geographic area. These results support that, on top of main human migrations, long ongoing socio-cultural factors have also shaped the genetic diversity observed at rural populations.

## Introduction

The current genetic variation of European human populations at a macro-geographic scale is mostly driven by isolation by distance patterns [1] and relatively modern massive migrations [2]. These results have been also observed at a country-based geographic scale [3–6]. Nevertheless, despite this general trend, some populations appear as genetic outliers, mostly related to geographically isolated rural and religious areas [7], as well as regions showing peculiar orographic events, such as islands [8–10]. Self-identified ethnicity (i.e. Romani people [8]) and ethnolinguistic background (such as Basques [11] can also promote genetic barriers at different scales.

A basic question is whether there are socio-cultural factors in Europe that remained immutable over time and affected mate preferences at a microgeographic scale. This question is complex, as the recent demography of European populations has been shaped by a progressive regional concentration [12]. This phenomenon is mainly explained by the continuous migration from rural areas towards industrialized urban areas during the last century [13]. Consequently, micro-geographic patterns can be expected to be currently blurred at urban areas, and patterns shaped by cultural factors present at rural areas obscured by their recent depopulation.

Studying a relatively isolated European rural geographic region showing geohistorical singularities that maintain over time political and cultural borders could help understanding the role of different socio-cultural factors in mating preferences during the last centuries. Within this context, the Catalan Pyrenees appear as a good candidate. First, the Catalan Pyrenees constitute a vast and abrupt region in the north of the Iberian Peninsula, delimited to the east by the Mediterranean Sea, to the west by the Aigüestortes National Park and the Noguera Ribagorçana river valley, to the north by the border with France and Andorra and to the south by the central depression of Catalonia. A marked orography is distributed on a west-east axis along the Prepyrenees massifs (see supplementary figure 1).

Most of the Pyrenean valleys run from north to south, except for the maritime ends and the Puigcerdà-La Seu d’Urgell axis. The Pyrenean valleys, of river origin, run in parallel to the watershed, with narrow gorges, which does not facilitate communications between valleys [14]. Along its entire extension, north-south communication axes are imposed, following the fluvial courses, which force its residents to move to the lower regions to ensure intra-Pyrenees relations. Furthermore, communications between valleys have been severely limited over time. It is at the end of the 19th and beginning of 20th century in which the construction of roads begins, first following the course of the rivers in a north-south axis, and much later the first road connections between the valleys were established. The limitations in the connections are strongly marked on the western slope, valleys of the Noguera Pallaresa and Segre rivers, while on the eastern slope, much less steep, the communications network is denser and earliest [15] (see supplementary figure 2).

The sum of all these mountain territories accounts for approximately a quarter of the surface of Catalonia, while the current population (70,000 people) barely exceeds 1% of the total Catalan population. Throughout history, the demography of the Pyrenean region has undergone marked variations. The times of maximum splendor go back to the Middle Ages, followed by successive demographic oscillations. In Modern times, the maximum population was reached in the 1880s, followed by a progressive population decline until today [16]. Due to the orography of the region, a distinctive feature of the Pyrenees settlements is its scattering, with villages mainly centered around the rivers and lower part of the valleys (see Figure 1).

**Figure 1.**
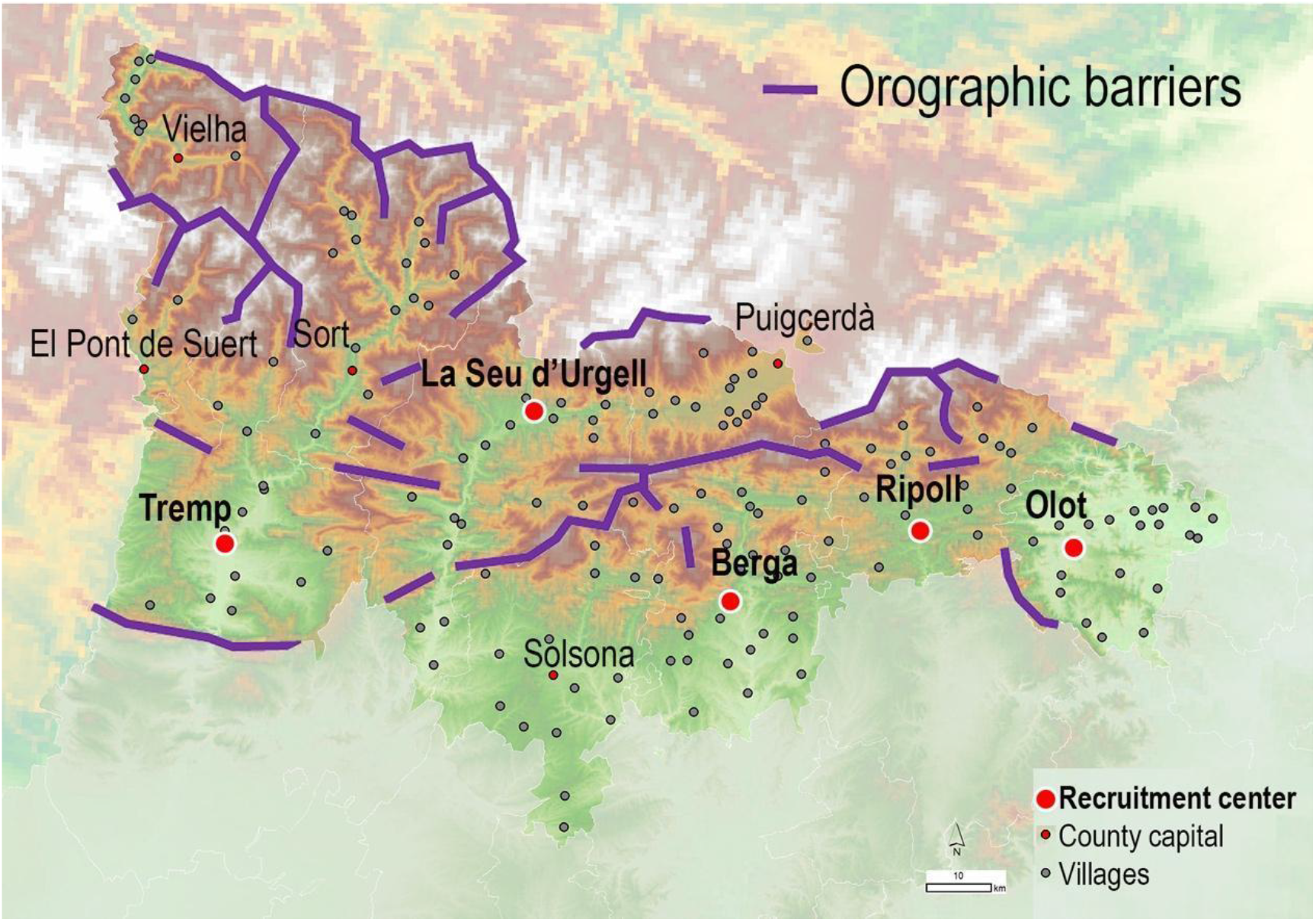
Orographic map of the Catalan Pyrenean region considered in this study, depicting the location of villages (gray dotted points), *comarca* capitals (red dotted points), and recruitment centers (white-red circles). Purple lines designate the main orographic barriers.

Finally, from a political point of view there are six Pyrenean *comarques* of Catalonia, corresponding to Alt Urgell, Alta Ribagorça, Val d’Aran, Pallars Jussà, Pallars Sobirà and Cerdanya and the northern part of another four more, Solsonès, Berguedà, Ripollès and Garrotxa (supplementary figure 3F). Administrative-social boundaries have remained stable practically since the Middle Ages. The oldest administrative divisions correspond to the limits established between bishoprics, with the bishopric of Urgell (Diœcesis Urgellensis) being the oldest (531 A.D.), followed by the Diocese of Vic (Diœcesis Vicensis, 561 AD), Girona (Diœcesis Gerundensis, 4th century), Lleida (Diœcesis Ilerdensis, 12^th^ century) and Solsona (Diœcesis Celsonensis, 16th century) (supplementary figure 3A). Other political-administrative divisions have been superimposed on these administrative divisions, such as the Middle-Age counties (11th century) (supplementary figure 3B), *Vegueries* (12th-18th centuries) (supplementary figure 3C), *Corregimientos* (18th century) (supplementary figure 3D), Provinces and *Comarques*, (19th century) (supplementary figure 3E) [17,18]. Despite these multiple denominations, the orographic backbone has imposed that they present great coincidences in their limits, so we could consider that these divisions have perpetuated the limits established from the old bishoprics with few variations (see supplementary figure 3).

In a previous study using whole genome sequencing on a limited sample size of 29 individuals [19], we showed that populations from the Catalan Pyrenees were interesting for understanding the genetic diversity of Spanish rural populations. More importantly, we observed the presence of micro-population substructure in this area, despite just covering populations in a region of around 200 km. Nevertheless, the lack of a continuous geographic sampling over the whole area of that study and the low sample size prevented identifying the atomic units of such substructure, as well as testing the factors that shaped the distribution of their genetic variation in the last millennia.

In the present study we have conducted a large genotyping study on 435 samples, analyzing and interpreting their genetic diversity in orographic and cultural terms.

## Methods

### Sample recruitment

We have collected samples from five reference hospital centers belonging to three health regions expanding the Catalan Pyrenees. Recruitment period was extended from July 2014 to October 2016. In total we recruited 435 samples (51.3% females) of subjects born and with all their grandparents born in the Pyrenean region (see Figure 1).

The data set represented an oldest extract of the population, with an average age of 55.6 years and equal proportion of sexes (table 1). Therefore, this sample should be considered moderately affected by the demographic changes that occurred during the 19th and 20th centuries. All subjects signed an informed consent, and the study had the approval of the Ethics Committee of the University of Barcelona.

**Table 1.**
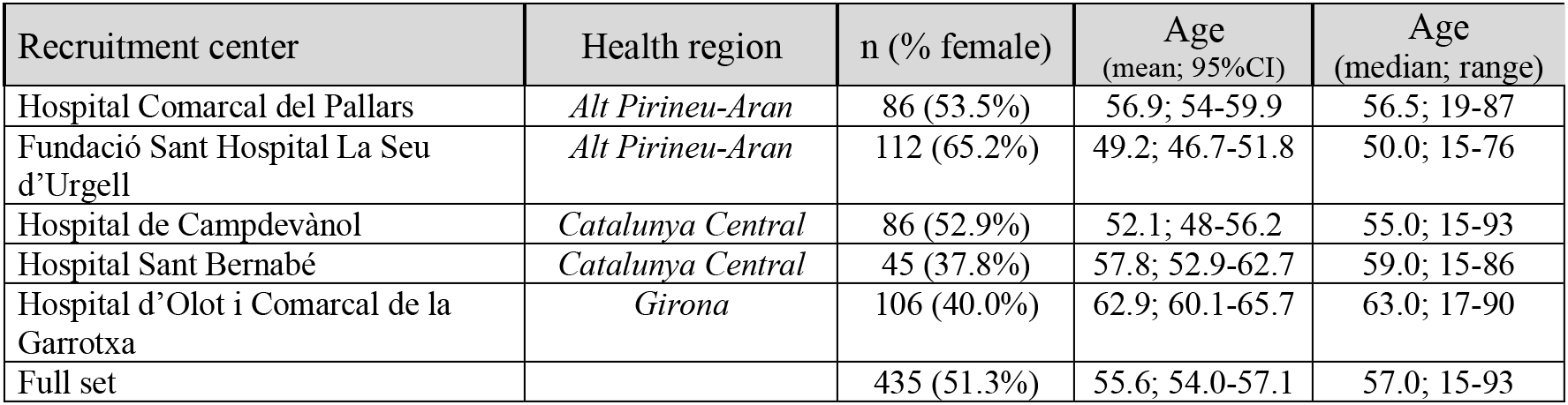
Distribution of samples according to recruitment center

For each participant we picked up demographic data such as sex, age and place of birth of the participant and place of birth their father and mother and for all their four grandparents. Birth places were assigned to geographic coordinates using *Vissir3*, the official viewer of the Cartographic and Geological Institute of Catalonia [20]. Assignment of centroid location of participants was done by calculating the mean longitude and latitude of grandparents (see supplementary table 1). Regional cartographic base maps and the terrain elevation model were from the Cartographic and Geological Institute of Catalonia. Maps were made with ArcGis 8.1 software (ESRI, Redlands, CA: Environmental Systems Research Institute), using facilities at the service of Cartography & SIG, of the University of Lleida (UdL).

### Genotyping

We collected 1ml of peripheral blood from each participant in EDTA-coated tubes. All participants were genotyped using Applied Biosystems Axiom™ Precision Medicine Diversity Array (PMDA) on GeneTitan™ Multi-Channel Instrument (Thermo Fisher, CA, USA) at *Centro Nacional de Genotipado* facilities (CEGEN, nodo Santiago de Compostela, Spain). Genotype clustering was analyzed using the Axiom Analysis Suite 6.0 software adjusting for BestWorkflow settings. Quality control (QC) was tuned to sample calling rate > 0.95, marker calling rate > 0.90, and Hardy-Weinberg equilibrium (HWE) > 10^−5^. After removing Indel and copy number variation (CNV), and QC, 780,310 SNPs (726,718 at autosomes, 49,319 at chromosome X, 406 at chromosome Y, 3457 at pseudoautosomal region and 410 at mitochondrial genome) on 435 subjects were available for subsequent analyses (GENPIR data set).

### Relatedness among individual participants

We estimated relatedness among samples by using SNPRelate R package [21] (genome-wide Identity by Descent (IBD_SNPRelate). IBD analysis reveals a degree of relatedness among several individuals (supplementary figure 4A). In order to select unrelated individuals for further analyses we selected one individual from each pair of individuals having a kinship ≥ 0.09 (supplementary figure 4B). This yields a final set of 397 unrelated individuals.

### Data Phasing and IBD tract estimation

Data was phased using the Michigan-imputation server-1.5.7 pipeline [22] with the eagle-2.4 phasing [23].

IBD tracts between pairs of chromosomes from different individuals and homozygote by descent (HBD) fragments between chromosomes from the same individual were inferred with Beagle 4.0 (IBD_Beagle) [24], using the human recombination map provided by the same software [24].

### Merging with reference populations

To make comparisons with other populations, GENPIR genotype data was merged with genotype data from samples of the 1000 Genomes Phase 3 project (1KG3) (2504 individuals) [25] and Human Genome Diversity Panel (HGDP) (929 individuals) [26]. After merging with the GENPIR dataset, a total of 3,868 individuals and 694,644 SNVs were selected with a genotyping rate of 0.998 (648,152 at autosomes, 46,315 at chromosome X, 177 at chromosome Y). We also applied unrelatedness criteria to the merged data set resulting in a final set of 3,721 unrelated individuals that were used for further analyses.

### Feature extraction analyses

#### PRINCIPAL COMPONENT ANALYSIS (PCA)

PCA was done on the unrelated set of GENPIR and merged datasets using plink1.9 software [27], after performing quality control (QC) that comprised: excluding markers with MAF<0.05 and pruning to *--indep-pairwise 200 25 0.3*. After QC passing, PCA analysis was done on 3,721 subjects and 175,277 autosomal markers. PCA QC control was also performed to tree additional data sets obtained after extracting subjects belonging to GENPIR and European populations (1048 samples; 137,907 autosomal SNVs); GENPIR and a subset of South European populations (871 subjects, 142,600 autosomal SNVs) and GENPIR population alone (397 subjects, 147,548 autosomal SNVs) in order to perform independent PCA analyses.

#### ADMIXTURE

ADMIXTURE v1.30 [28] was used to analyze the population structure and calculate the corresponding cross-validation error value (CV) of the considered clusters (K = 2–4). ADMIXTURE analysis was done on merged GENPIR and European populations (1048 samples; 137,907 autosomal SNVs) applying the same QC control as for PCA analysis. Distribution of derived European components from K=3, classified as Northeast (NE), Southeast (SE) and Southwest (SW), were compared among Pyrenean regions (West, Mid, East) by pairwise t-test.

#### MULTIPLE CORRESPONDENCE ANALYSIS

The bishopric origin of the grandparents from a given sample was projected in two dimensions using a multiple correspondence analysis [29] as implemented in the package Factomine [30] in R software [31].

#### SMACOF

Given the regionality of the sampled dataset, the presence of population substructure must be relatively small. Since the identical by descent (IBD) statistic is sensitive towards subtle hidden genetic relationships between individuals, we computed an IBD matrix with Beagle [24]. We projected the IBD_Beagle matrix in a lower dimensional space after transforming it into dissimilarity by means of a non-metric ordinal SMACOF analysis [32] implemented in R software.

#### ESTIMATING ANCESTRY PROPORTIONS ACROSS GEOGRAPHIC SPACE

We evaluated genetic ancestry across geographic space on GENPIR population autosomal data set by tess3r software [33], applying the following parameters: method = “projected.ls”, ploidy = 2, max.iteration = 200, rep = 10, keep = “best”, tolerance = 1e−05. We kept the most highly supported model (i.e., “best” based on cross-entropy scores) within each of the 10 replicates. Two subjects were excluded lacking centroid assignment. We also exclude markers with MAF<0.05 as well as pruning to --indep-pairwise 200 25 0.3.

#### MANTEL TEST & PROCRUSTES

The relationship between genetic variation and cultural or political units while controlling by the putative confounder effect of geography was tested by means of a partial Mantel test [34] implemented in the R package vegan [35].

A Procrustes analysis [36] between geography and the coordinates estimated by PCA or SMACOF using genetic variation was computed using the R package vegan.

#### MCLUST

Unsupervised clustering of the individuals using the first two dimensions computed by SMACOF was conducted by means of the MCLUST package [37] implemented in R. Mclust models the points as a finite mixture of Gaussian distributions. Ascertainment of the best number of clusters was conducted based on BIC metrics.

## Results and discussion

### The genetic diversity of the Catalan Pyrenees in the context of worldwide populations

We first analyzed the genetic relationships of the sampled individuals at worldwide level using a Principal Component Analysis (PCA) (supplementary figure 5).

The first two PCA components (PCs), explaining 11.3% of variance, show that the considered Catalan Pyrenees samples are genetically related to other samples with European ancestry (supplementary figure 5A). Within Europe, the Catalan Pyrenees samples fall within the first two PCs (explaining 0.7%) close to samples from Southern Europe and using Southern Europe samples (first two PCs explaining 0.5%), close to individuals from Iberian and French Basque origin (supplementary figure 5B & C).

Given these results, we run an unsupervised ADMIXTURE analysis focusing on the European region for K ranging from one to eight in order to quantify the European ancestry proportions in GENPIR samples, revealing a minimum cross validation error value for K= 2 - 3 (supplementary figure 6).

Considering the three major components in the European region revealed at K=3 (geographically classified as “Northeast (NE)”, “Southeast (SE)” and “Southwest (SW)”) (figure 2A), we estimated the proportion of these three components in our samples. Figure 2B&C shows a decreasing gradient for the contribution of NE and SE components being highest at the East, intermediate at the Mid and lowest at the West Pyrenean regions. In contrast, the SW European component shows an increasing gradient being lowest at the East, intermediate at the Mid and highest at the West Pyrenean regions. These results agree with the longitudinal gradient observed on the same region by other studies using classical markers [38], as well as with the main axes of genetic differentiation within the Iberian Peninsula [3]. Therefore, all these results support that the Catalan Pyrenees fall within the genetic continuum within the Iberian Peninsula. At a local level, the presence of a higher proportion of SW European ancestry in the West region of the Catalan Pyrenees supports that this region has been more accessible for migration than other populations from the Iberian Peninsula. In addition, the higher contribution of NE and SE European components at the East region of the Catalan Pyrenees reflects a major permeability of this Pyrenean region, where geographical barriers are much weaker and with a much richer communications network, that has allowed contact with the nearby regions of northern and eastern Europe.

**Figure 2.**
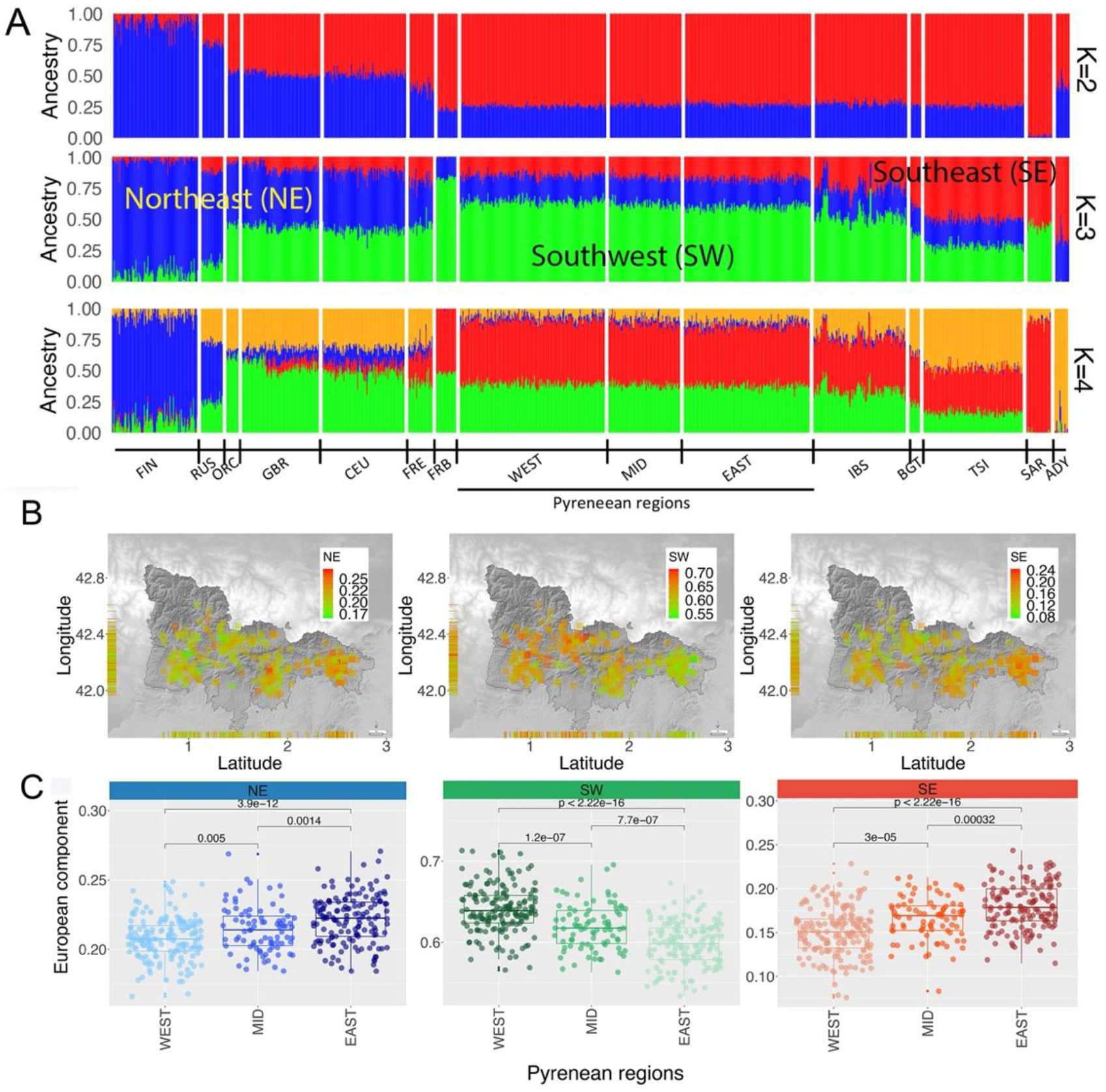
Admixture analysis of European populations (FIN, Finland; RUS, Russia; ORC, Orcadian (Scotland); GBR, Great Britain and Scotland; CEU, Utah residents (CEPH); FRE, French; FRB, French Basques (France); IBS, Iberian Peninsula (Spain); BGT, Bergamo (Italy); TSI, Toscana (Italy); SAR, Sardinia (Italy) and ADY, Adygei populations) and Catalan Pyrenean populations (West, Mid and East regions) for K=2 to K=4 (A). For K=3 we depicted Northeast (NE) European component (FIN, RUS); Southwest (SW) European component (FRB, IBS) and Southeast (SE) European component (BGT, TSI, SAR). B) Distribution of K=3 European components (NE, SW and SE) on GENPIR individual samples according to their geographical location. C) Box plot distribution of K=3 European components (NE, SW and SE) in the West, Mid and East Catalan Pyrenean regions, including P-values of pairwise t-test comparisons.

### Population substructure is present at a microscale level within the Catalan Pyrenees

Given the observed differences in ancestry among the GENPIR samples, we conducted a PCA only using GENPIR samples (figure 3; 0.5 % of variance).

**Figure 3.**
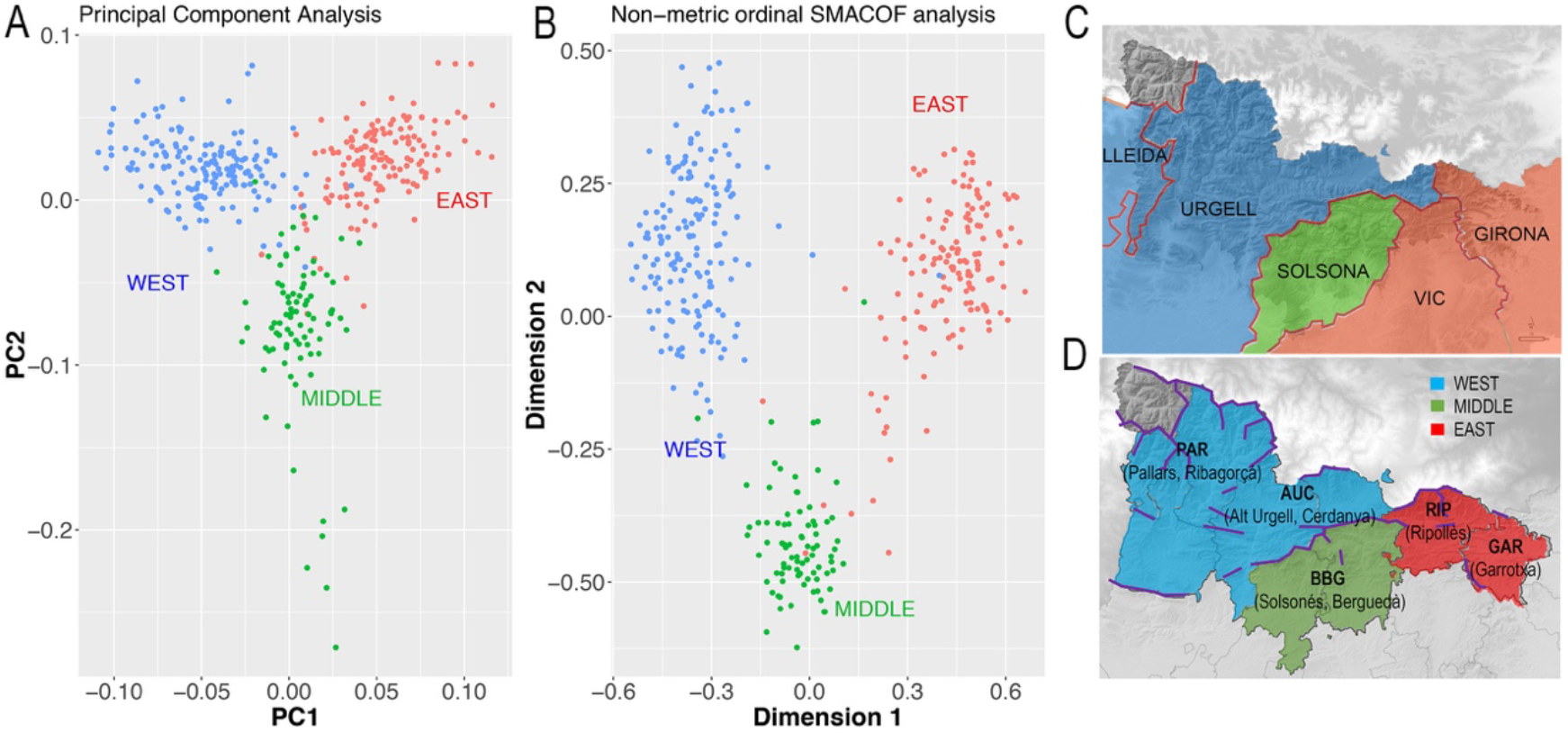
A) First two PCs, explaining 0.5% of variance of a PCA of GENPIR individuals. B) First two dimensions out of the four-set obtained by projecting an IBD matrix between pairs of individuals using an ordinal SMACOF analysis. GENPIR samples are assigned to three Catalan Pyrenees regions according to their centroid assignment, C) Map denoting assignment of bishoprics at each region: WEST (blue, bishoprics of Lleida and Urgell); MIDDLE, (green, bishopric of Solsona); EAST (red, bishoprics of Vic and Girona). D) Map denoting assignment of *comarques* at each region: WEST (blue, *comarques* of Pallars, Ribagorça, Alt Urgell and Cerdanya); MIDDLE, (green, *comarques* of Solsonès and Berguedà); EAST (red, *comarques* of Ripollès and Garrotxa). In D, the purple line denotes main orographic barriers.

We observed that the genetic map defined by the two PCs supports the presence of a non-random distribution of GENPIR individuals. Individuals from West, East and Middle Pyrenean regions tend to be grouped into three prominent clusters (figure 3A). Since shared haplotype tracts are more sensitive for identifying subtle population substructure [39], we wondered if the pattern observed by the PCA could be enhanced by projecting the IBD matrix between pairs of individuals estimated from Beagle 4.0 [24]. We applied an ordinal SMACOF approach, in which the distance between pairs of individuals is interpreted in rank terms rather than by its magnitude. This decision was taken due to the fact that the overall genetic differences between all individuals are expected to be limited. Therefore, the presence of few outliers showing a higher degree of relationship (even after excluding highly related individuals) could distort the projection towards these relationships rather than describing the patterns observed among all samples. The first two dimensions from the SMACOF analysis considering four dimensions (stress = 0.4) produce a similar picture as observed by PCA (figure 3B). However, it enhances the differences between the three different clusters of populations, supporting micro substructure in the Pyrenees.

Furthermore, this micro-population substructure can be interpreted in geographic terms. In particular, a procrustes analysis between the geographic coordinates of the samples and the genetic map defined by the first two PCA dimensions showed a symmetric correlation of 0.61 (p-value after 10,000 permutations < 9.999e-05).

Taking into account the orography of the region, the Cadi Massif constitutes a clear geographic barrier detaching the Western to the Middle and Eastern regions while no clear geographic barrier appears to detach the Middle to the Eastern regions (supplementary figure 1). When applying actual (*comarques*) and historical (bishoprics) administrative-religious boundaries, the West cluster comprises Pallars, Ribagorça, Cerdanya and Alt Urgell *comarques* as well as Lleida and Urgell bishoprics; the Middle cluster comprises Solsonès and Berguedà *comarques* and Solsona bishopric and the East cluster includes Ripollès and Garrotxa *comarques* as well as Vic and Girona bishoprics (figure 3 C & D).

### Micro-population substructure within the Catalan Pyrenees is better explained by geohistorical organization (religious) structures

Given these results supporting a geographic interpretation of the genetic variation, we wondered whether models incorporating geographic information could identify genetic barriers present between the samples. The ancestry map generated by tess3 at K=2 to K=4 (figure 4) identifies a strong longitudinal component that agrees with ADMIXTURE analyses and procrustes, and splits individuals following a geographic partition that matches the genetic differentiation observed in previous analyses.

**Figure 4.**
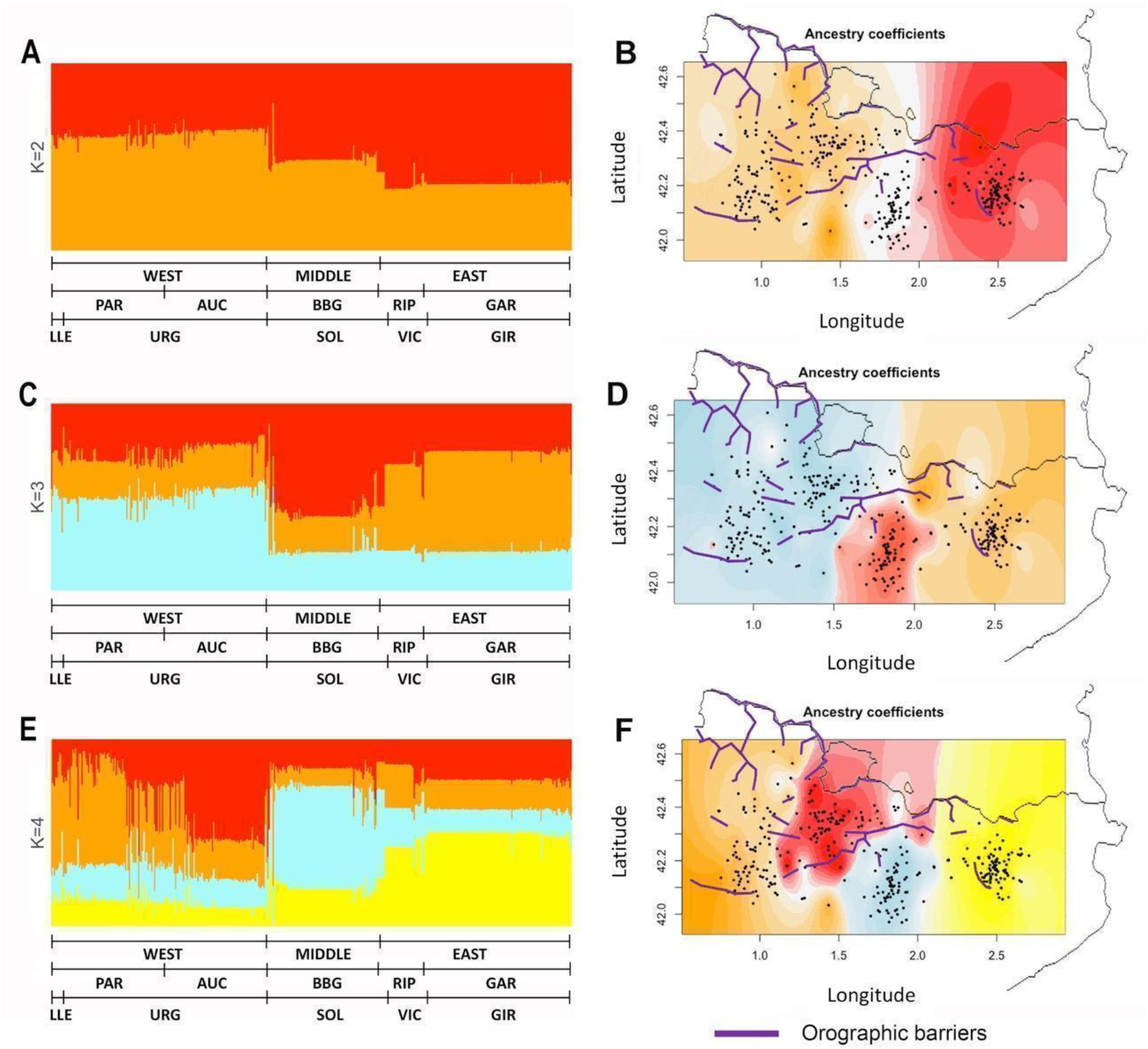
Spatial distribution analysis of the ancestry of GENPIR individuals. Barplot (A, C, E) and Geographical (B, D, F) pattern distribution of ancestry components of GENPIR samples assuming K=2 to K=4. In A, C, E GENPIR samples are assigned to WEST, MIDDLE or EAST Catalan Pyrenees regions, *comarques* (PAR, Pallars-Ribagorça; AUC, Alt Urgell-Cerdanya; BBG, Berguedà and Solsonès; RIP, Ripollès and GAR, Garrotxa) and bishoprics (LLE, Lleida; URG, Urgell; SOL, Solsona; VIC, Vic and GIR, Girona), according to their centroid assignment. In B, D, F individual values of the ancestry components of GENPIR samples for K= 2 to 4 are plotted over the geographic location of each subject. Purple line denotes main orographic barriers.

Interestingly, these divisions do not fully agree on orographic events, such as the presence of mountains, that could act as genetic barriers. Particularly, the division that is observed between East and West at K=2 is set in a region lacking clear geographic barriers. Therefore, as previously considered, other factors correlating with geography must be shaping the observed genetic differentiation across the region. We also point out that at K=3 the algorithm mainly reproduces the West, Middle and East clustering, which we previously observed with both PCA and SMACOF analysis.

We wondered if such genetic clusters identified in previous analyses were shaped by current administrative divisions (*comarques*), or if they were influenced by other long term ongoing cultural factors such as the limits defined by middle age-founded bishoprics. We applied a Mclust analysis using the first two dimensions from SMACOF in order to identify clusters showing a similar probability distribution function. The optimal number of clusters identified by Mclust was six (BIC = 406.33) that accurately reflect a hardest effect of geographic barriers that is shaped by administrative divisions, both *comarques* and bishopric (figure 5). Cluster I includes subjects from the *comarques* of Pallars and bishopric of Lleida, cluster II subjects from the *comarques* of Pallars and Alt Urgell as well as bishopric of Urgell, cluster III mainly includes subjects from the comarca of Alt Urgell and bishopric of Urgell, cluster IV subjects from *comarca* of Solsones and Berguedà as well as bishopric of Solsona, cluster V subjects from *comarca* of Ripollès and bishopric of Vic and cluster VI mainly includes subjects from *comarca* of Garrotxa and bishopric of Girona (figure 5).

**Figure 5.**
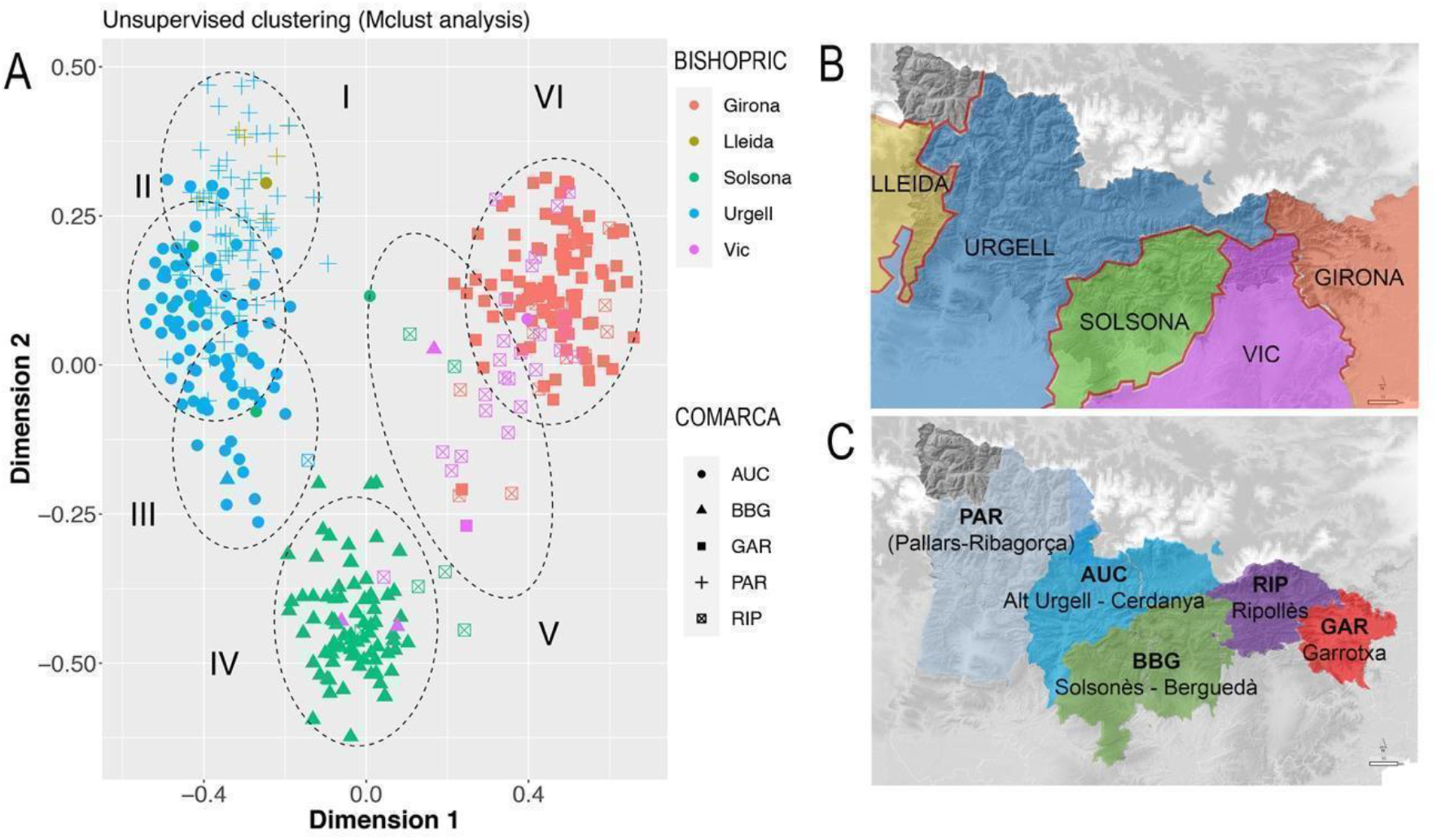
A) Suggested clusters (ellipses I to VI) identified by applying the unsupervised clustering Mclust analysis. Colors identify bishoprics (Girona, red; Lleida, brown; Solsona, green; Urgell, blue and Vic, purple). Shape identify *comarques* (circles AUC, Alt Urgell-Cerdanya; triangles BBG, Berguedà-Solsonès; squares GAR, Garrotxa; crosses PAR, Pallars Jussà-Pallars-Sobirà-Ribagorça; crosses in squares RIP, Ripollès). B) Colored bishoprics according to clustering, colors as in A. C) Colored *comarques* according to clustering, (light-blue PAR, Pallars Jussà-Pallars-Sobirà-Ribagorça; blue AUC, Alt Urgell-Cerdanya; green BBG, Berguedà-Solsonès; purple RIP, Ripollès and red GAR, Garrotxa).

Interestingly, such bishopric division from the SMACOF bi-plot is also observed within the geographic region (depicting ~70 km far away) that contains the intersection of four bishoprics (see supplementary figure 7).

Since bishoprics and *comarques* depend on geography, the identified geographic clusters by tess3 could reflect cultural and/or political barriers. We conducted a partial Mantel test between the Euclidean distance estimated from the first two PCs, and the assignment to the same administrative division or bishoprics, controlled by the geographic distance. Pearson’s correlation between PC based distances and bishoprics after controlling by geography in a partial Mantel test was 0.434 (p-value <1e-04 using 9,999 permutations). This degree of association is higher than the observed between genetics and administrative divisions of *comarques* (Pearson’s correlation = 0.324, p-value <1e-04 using 9,999 permutations). When considering a classification based on bishoprics and administrative divisions at the same time, Pearson’s correlation decreased to 0.294 (p-value <1e-04 using 9,999 permutations). Similar results supporting higher correlation between bishopric divisions and genetic diversity were obtained when using the first two dimensions from the SMACOF analysis conducted with the IBD distance matrix. The degree of correlation between genetics and bishoprics after controlling by geographic distance was 0.563 (p-value <1e-04 using 9,999 permutations). This correlation decreases to 0.382 (p-value <1e-04 using 9,999 permutations) when considering administrative divisions and it is similar when considering both bishopric and administrative divisions (R = 0.361, p-value <1e-04 using 9,999 permutations). Overall, these results suggest that bishoprics play a role in shaping the genetic variation of GENPIR samples by modifying mating preferences.

We supported such a hypothesis based on genetic data by taking advantage of the sampling strategy conducted in this study, which incorporated the geographic birthplace of the four grandparents (Figure 6, supplementary table S1).

**Figure 6.**
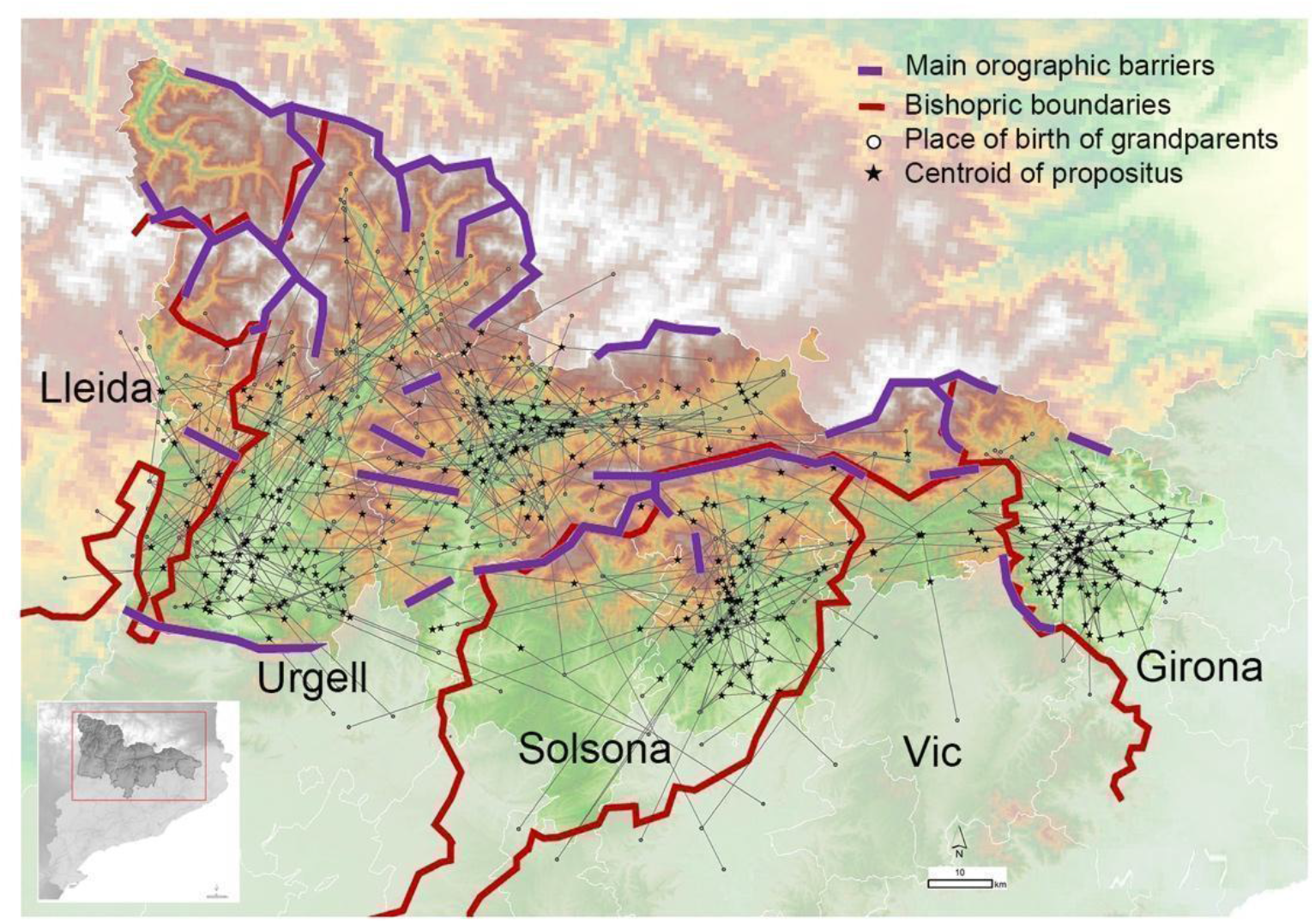
Geographic location of the individuals with regards to the mating relationship of grandparents. Each *propositus* is linked to their grandparents generating a complex network centered on bishoprics. Red lines delimit the bishopric boundaries. Purple lines indicate the presence of orographic events preventing direct routes of contact between regions.

figure 6 shows the grandparent to *propositus* network in which each propositus is linked to their four grandparents generating a complex network centered on bishopric boundaries. We can also observe that grandparent and parent couples come from the same bishopric (over 80% of couples) or to a nearest bishopric (near 10% of couples) (figure 7A). The overall relationship between bishopric and mating was summarized by a multiple correspondence analysis (MCA) using the bishopric of each grandparent, parent and the considered sample. The relationship defined by the bishopric origin of the grandparents and samples is summarized by a triangle in the first two dimensions (explaining 39.3% of total inertia; figure 7B) of the MCA. The first vertex contained relationships from the bishoprics of Urgell and Lleida. The second vertex was determined by the relationship between members from the bishopric of Solsona and the third one by the bishopric of Girona. The grandparents and current samples from the bishopric of Vic fall within the axis defined by Solsona and Girona, indicating a higher number of bishopric-admixed grandparent couples with the bishoprics of Solsona and Girona (see figure 7B). The center of the triangle depicted relationships where at least one of the grandparents is out of the bishoprics. Overall, the observed mating relationships agree with the geographic proximity between bishoprics (Lleida and Urgell, Vic between Solsona and Girona), as expected given the dependence between geography and these geohistorical organizations, and highlight the bishopric-based mating choice of the individuals from GENPIR.

**Figure 7.**
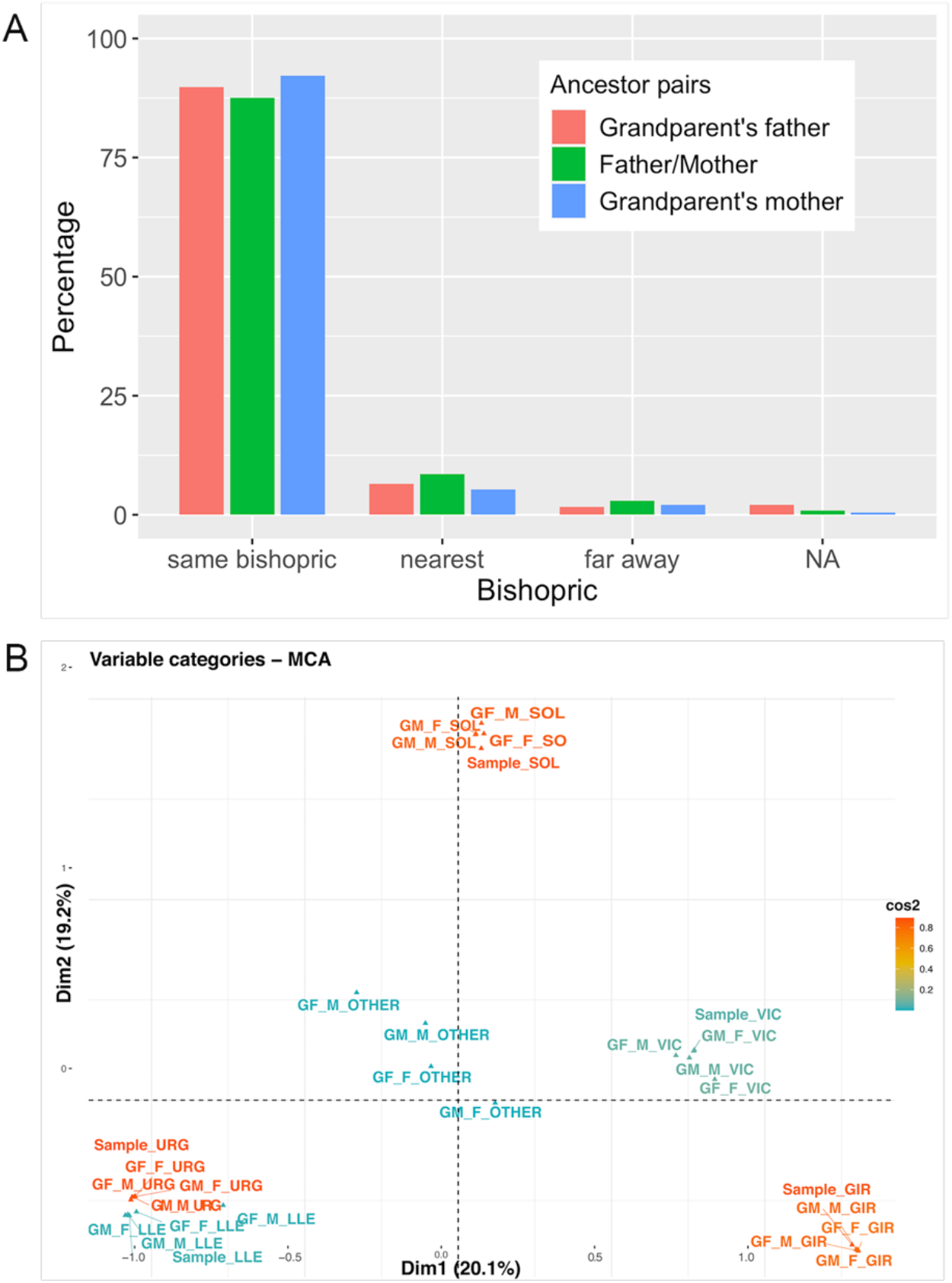
A) Relationship of marriages depending on the bishoprics of parents and grandparents couples. B) Multiple Correspondence Analysis between the bishopric of the sample and her/his parents and grandparents.

The ecclesiastical division of the modern era responds to human spheres of multisecular relationships. For this reason, it is not surprising that the bundles of family relationships roughly coincide with the bishoprics: a central Pyrenean space ‒bishopric of Urgell‒ with abundant interactions (although it is capable of being subdivided into *comarques*); an eastern area (Girona); and a southern territory or central Catalonia (bishopric of Solsona, perhaps undifferentiated from that of Vic).

Traditionally, the main force used for explaining biases in mating choice in Spanish rural areas compared to urban areas has been geography [40]. Poor transportation facilities and limited communications-which in turn depend on orography- enhance the practice of consanguineous marriages and inbreeding in Spain. In particular, in small rural and dispersed localities, a limited number of suitable local partners conditioned marriage, and marrying a distant relative was a likely option when few other partners were available. This situation was even more common when the population started increasing due to medical advances, but mobility remained restricted [41]. Furthermore, in a sociocultural context, aunt-nephew and uncle-niece marriages could have been practiced in rural areas in order to maintain or enlarge the family inheritance [40]. Similarly, the regional practice in Northern and Northwestern Spain from the 18th to the 20th century of a complex household system could have favored first cousin mating in these rural areas [40]. Overall, these socio-cultural processes could explain the overall patterns of isolation by distance that we detected in our different analyses. However, our study also supports that bishopric can act on shaping the genetic variation of rural areas. This observation can be interpreted in the socio-cultural context of the Catalan Pyrenees. In this region, the old urban centers of cities and towns with episcopal seats are the first location of markets or fairs. Cities such as La Seu d’Urgell (bishopric of Urgell), Besalú (bishopric of Girona), Berga (bishopric of Solsona) or Vic (bishopric of Vic) have documented markets or fairs since XII and XIII centuries. In rural areas, markets and fairs act as a space for sociability promoting personal contacts between relatives and acquaintances. Social relationships were established or renewed, labor contracts were established and marriages were arranged [42].

## Conclusions

Humans, as social animals, are subject to rules that emanate from power, economic, political or religious, which beyond allowing the proper functioning of social structures, also modulate interpersonal relationships. Throughout history the role of the church has been prominent, acting as a central and determining factor of interpersonal relationships in past societies. In particular, the Catholic Church, with its rigid pyramidal structure, has left an imprint on societies that go beyond religious aspects. As our results show, the genetic structure of the populations of the Catalan Pyrenees, in addition to the geographical factor, has also been modulated by administrative-social factors, which have established limits to the social relationships of individuals, being the limits established by the bishoprics one of the determining factors.

In this context, the bishoprics can be considered as the most representative territorial organization frameworks, historically very stable and cohesive, and necessarily well adapted to the strong geographical conditions that mark the Pyrenean environment.

A singular orography that has determined the establishment of administrative-religious limits, perpetuated unalterably for centuries, and the rigid social structure enforced by the Catholic Church over long periods of time, have shaped the genetic structure of the Catalan Pyrenees population. It remains to be determined to what extent this structure may have generated interpopulation differences able to be reflected in differences on genetic susceptibility to diseases.

## Data accessibility

All data were included in the main figures and the accompanying electronic supplementary material, supplementary figures 1–7, supplementary table 1.

## AUTHORS’ CONTRIBUTIONS

J.F(1).: conceptualization, formal analysis, investigation, funding acquisition, methodology, validation, visualization, project administration, supervision, writing— original draft, writing—review and editing; I.M.: formal analysis; investigation; methodology; software; visualization; writing – review & editing; M.L.: formal analysis; investigation; methodology; software; visualization; writing – review & editing; M.G.: formal analysis; investigation; methodology; software; visualization; writing – review & editing; M.M.A.: formal analysis; investigation; methodology; visualization; writing – review & editing; J.B.: investigation; methodology; visualization; writing – review & editing; A.C.: investigation; methodology; visualization; writing – review & editing; J.F.(6): investigation; methodology; visualization; writing – review & editing; J.F.(7): investigation; methodology; visualization; writing – review & editing; J.G.: investigation; methodology; visualization; writing – review & editing; J.L.R.: investigation; methodology; visualization; writing – review & editing; J.S.: investigation; methodology; visualization; writing – review & editing; P.M.: conceptualization, funding acquisition, investigation, methodology, visualization, writing – review & editing; O.L.: conceptualization, formal analysis, investigation, methodology, validation, visualization, project administration, supervision, writing—original draft, writing—review and editing. All authors gave final approval for publication and agreed to be held accountable for the work performed therein.

## Acknowledgements

We thank all the volunteers who donated DNA samples and who were involved in the collection of samples and genealogical data (Hospital Sant Bernabé, Berguedà; Fundació Sant Hospital, Alt Urgell; Hospital d’Olot i Comarcal, Garrotxa; Hospital de Campdevànol, Ripollès and Hospital Comarcal, Pallars).

