## Supplementary figures 1-7 for "The power of geohistorical boundaries for modeling the genetic background of human populations: the case of the rural Catalan Pyrenees"

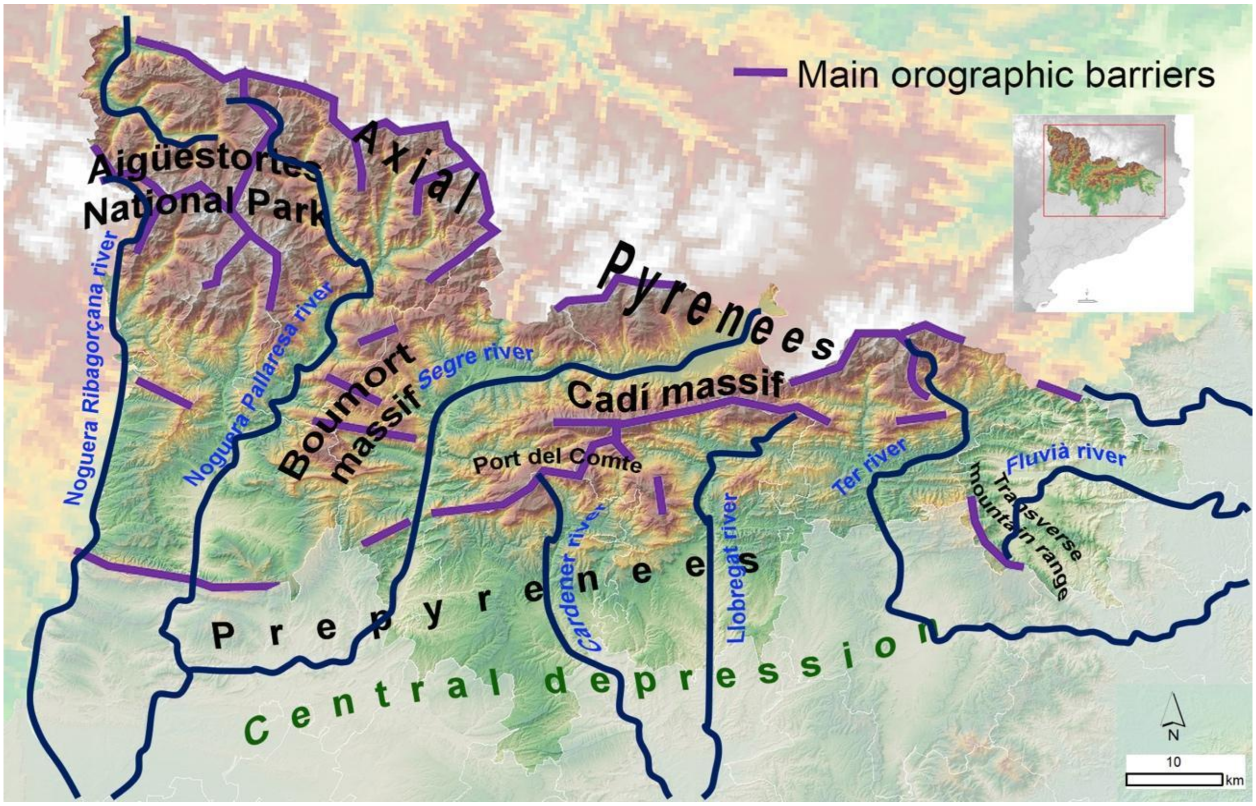


**Supplementary Figure 1**. Orographic map of the Catalan Pyrenean region considered in this study, depicting the most relevant orographic elements as massifs, rivers and depressions. Labeled by purple line the main orographic barriers defined by mountains higher than 1500 meters.

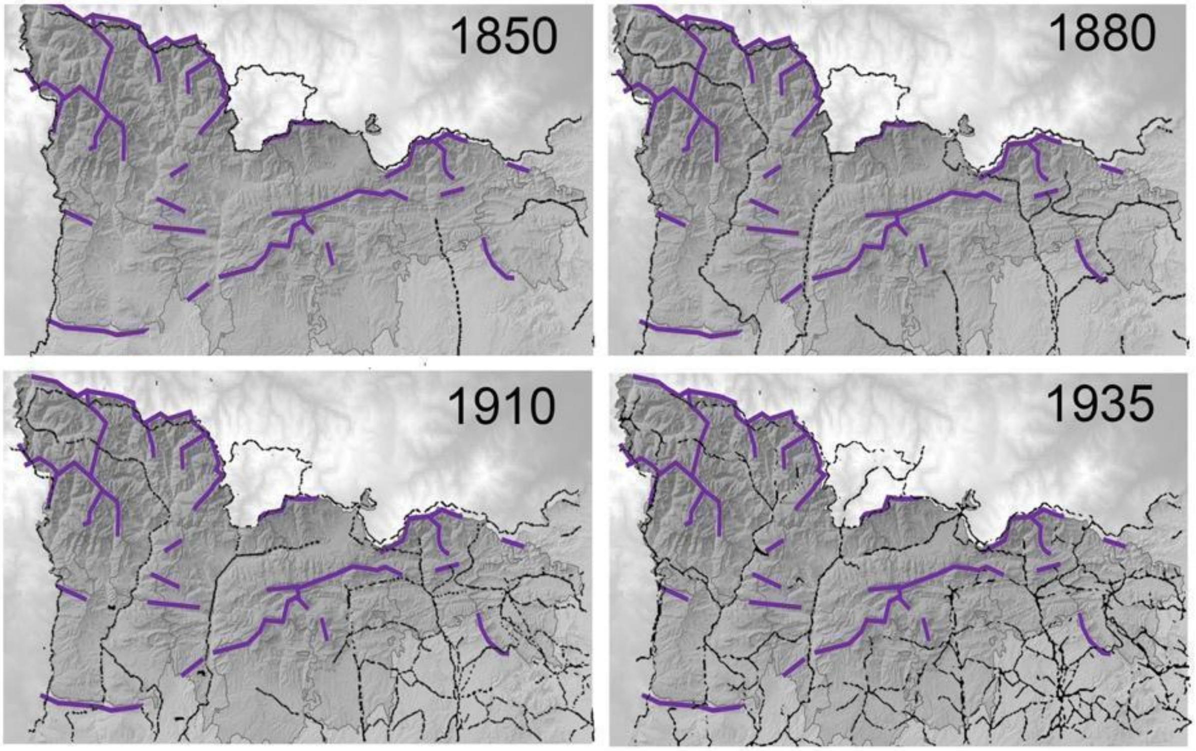


**Supplementary Figure 2.** Evolution of road communications in the Catalan Pyrenees over the last two centuries. Black dotted lines designate the growing road network from 1850 to 1935. Purple lines designate the main orographic barriers defined by mountains higher than 1500 meters. Modified from [15].

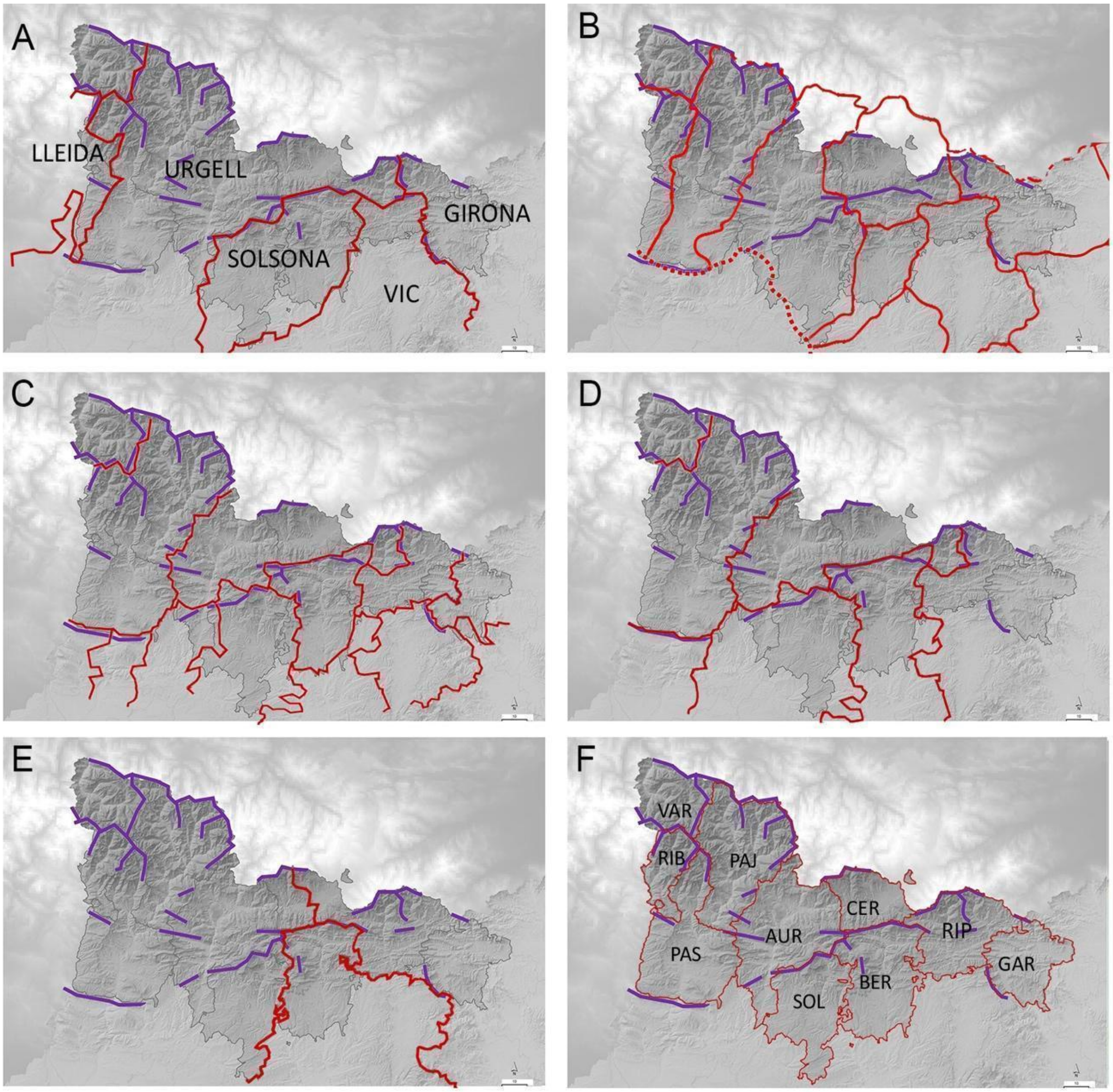


**Supplementary Figure 3.** Historical administrative divisions (A to F) at the Catalan Pyrenees over last centuries. A) Bishoprics boundaries dated on 531 AD to s.XVI. B) Middle-Age County boundaries. C) *Vegueries* boundaries at $12^{th}$ to $18^{th}$ centuries. D) *Corregimientos* at $18^{th}$ century. E) Provinces from $19^{th}$ century. F) Actual "*comarques*" and Val d'Aran. AUR, Alt Urgell; BER, Berguedà; CER, Cerdanya; GAR, Garrotxa; PAJ Pallars Jussà; PAS; Pallars Sobirà; RIB; Alta Ribagorça; SOL; Solsona; VAR Val d'Aran. Red lines designate administrative-religious borders. Purple lines designate the main orographic barriers. Adapted from [17,18].

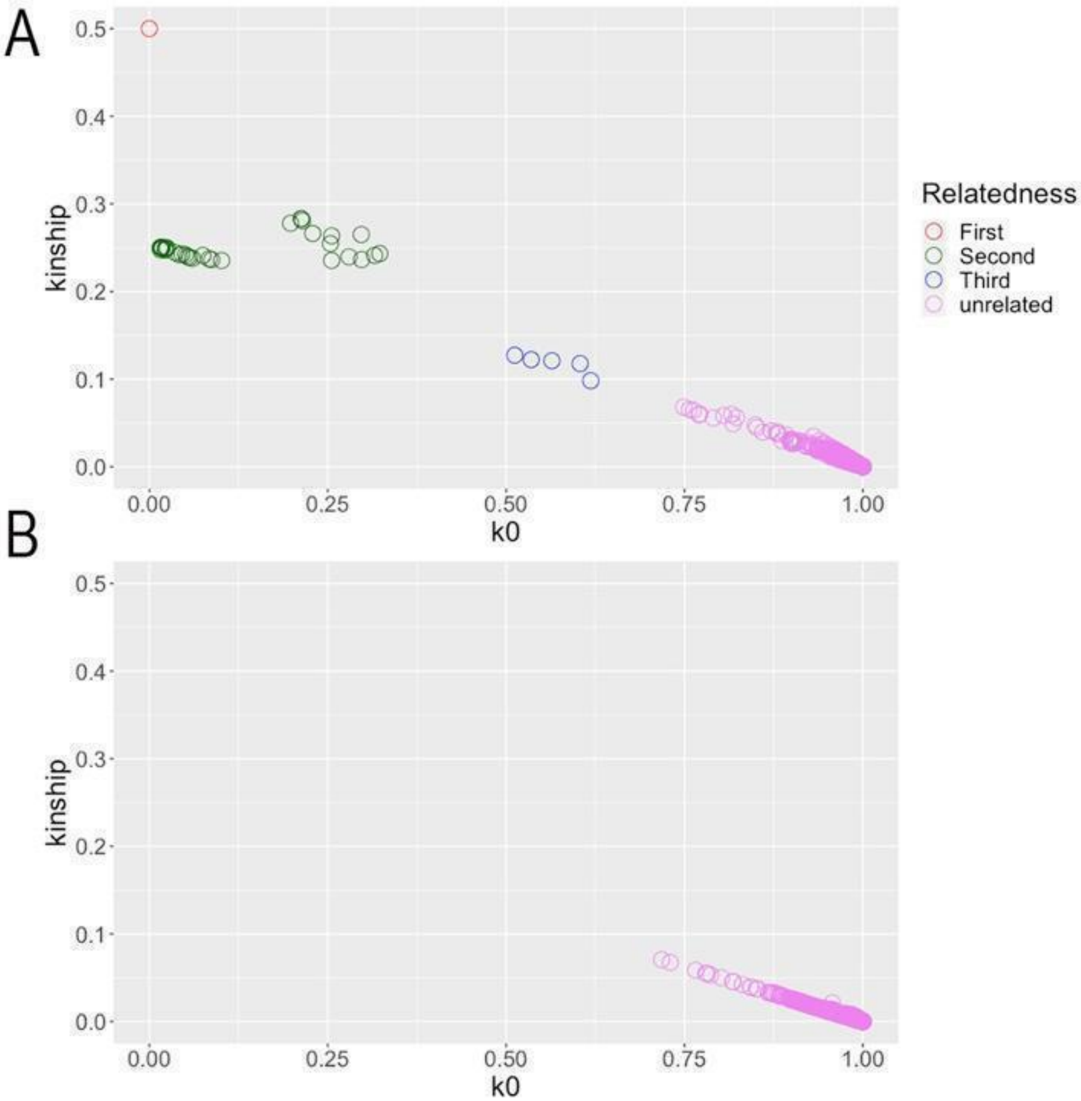


**Supplementary Figure 4**. Relatedness among individual participants. IBD analysis showing k0 vs. kinship values distribution in all 435 genotyped individuals (A), and after excluding those with kinship ≥ 0.09 (B). First degree relatives, (kinship 0.5; k0=0, red circles); second degree relatives (kinship~0.25; k0~0.25, green circles), third degree relatives (kinship~0.25 k0~0.5; blue circles) and unrelated individuals (kinship≤0.09; k0>0.7; pink circles).

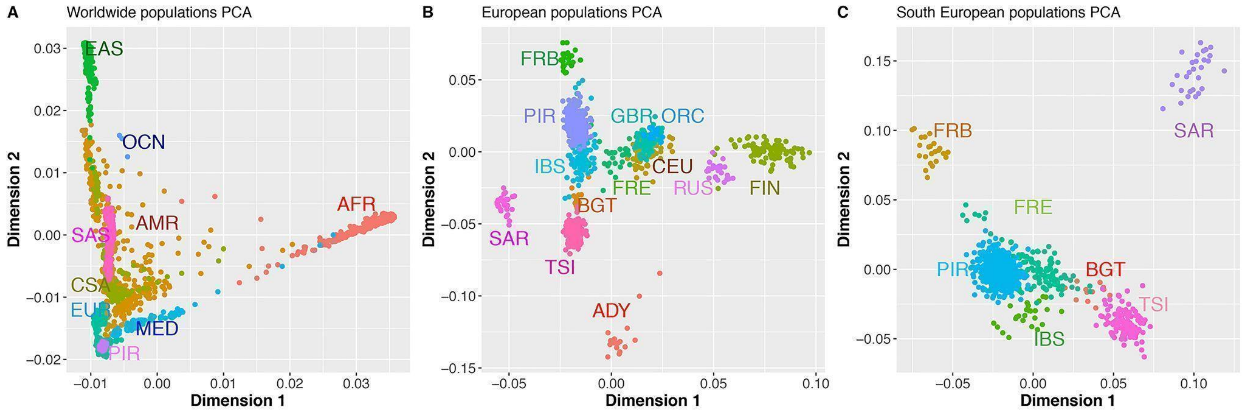


**Supplementary Figure 5**. First two PCs of PCA analyses of worldwide populations (A), European populations (B) and South European populations (C) including the GENPIR population. A) PCA analysis of world-wide superpopulations AFR, African; AMR, Amerindians, CSA, central Asia; EUR, Europeans; EAS, east Asia; MED, Middle east; SAS, South Asia and Catalan Pyrenean (PIR) samples. B) PCA analysis of European populations ADY, Adygei; BGT, Bergamo (Italy); CEU, Utah residents (CEPH) with Northern and Western European ancestry; FRB, French Basques (France); FRE, France; FIN, Finland; GBR, Great Britain and Scotland; IBS, Iberian Peninsula (Spain); ORC, Orcadian (Scotland); RUS, Russia; SAR, Sardinia; TSI, Toscana (Italy) and PIR, Catalan Pyrenean samples. C) Detailed PCA analysis of South European populations.

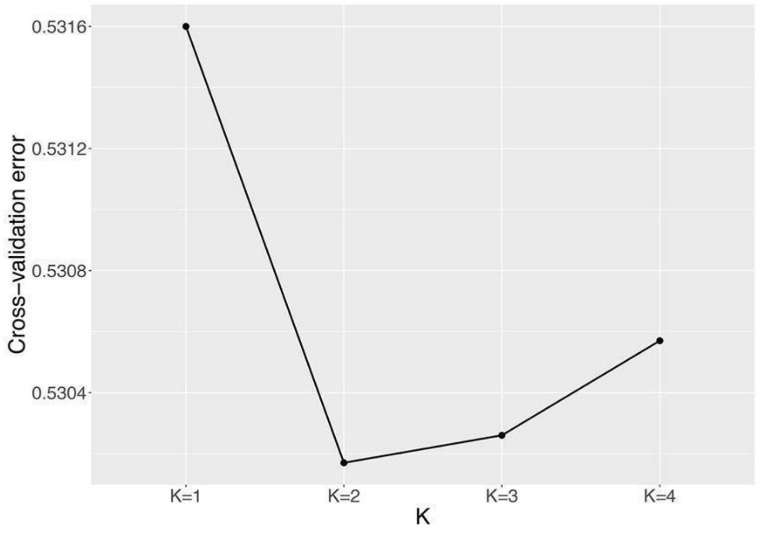


**Supplementary Figure 6**. Cross-validation errors (CV) of ADMIXTURE analysis for K=1 to K=4.

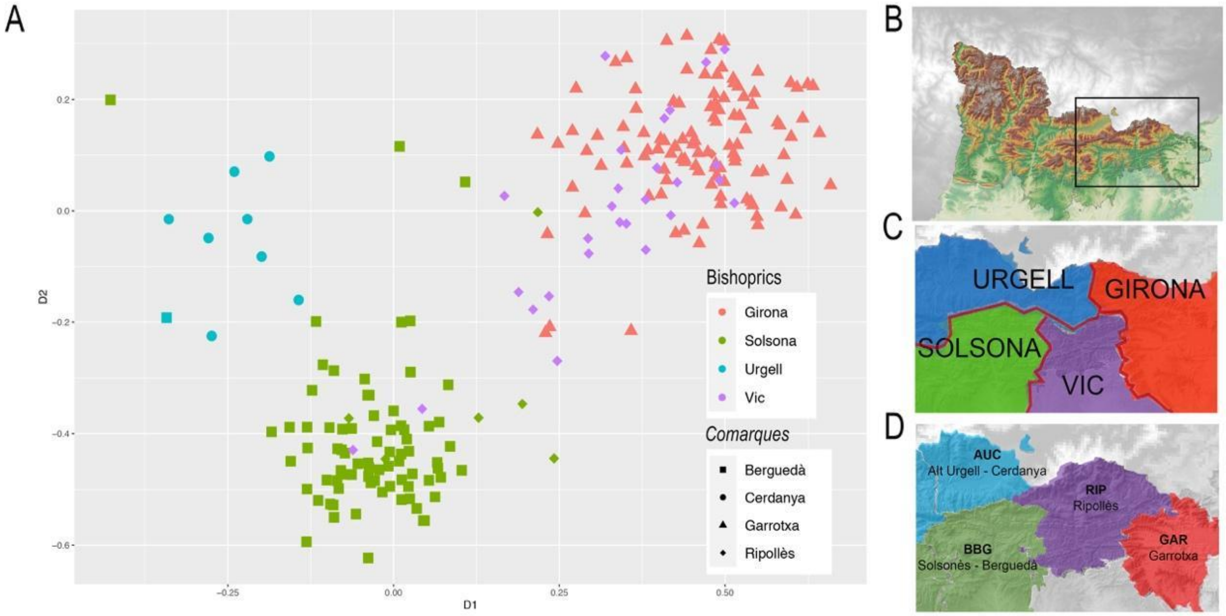


**Supplementary Figure 7**. A) First two dimensions of the SMACOF analysis depicted in figure 5 focusing on individuals (N=230) from the geographic region that contains the intersection of four bishoprics. B) detailed orography of the focused region. Despite the small distance considered, the influence of bishoprics (C) and *comarques* (D) is observed. In A, Colors identify bishoprics (Urgell, blue; Solsona, green; Vic, purple and Girona, red). Shape identify *comarques* (circles AUC, Alt Urgell-Cerdanya; square BBG, Berguedà-Solsonès; triangles GAR, Garrotxa and diamond RIP, Ripollès). B) Orography of the detailed region. C) Colored bishoprics according to clustering, colors as in A. D) Colored *comarques* according to clustering, (AUC, Alt Urgell-Cerdanya, blue; BBG, Berguedà-Solsonès, green; RIP, Ripollès, purple; GAR, Garrotxa, red).
