## Supplementary table 1 for "The power of geohistorical boundaries for modeling the genetic background of human populations: the case of the rural Catalan Pyrenees"

| DNA_code | gender | age_at_recruitment | Recruitment_place | Recruitment_center | Recruitment_comarca | Propositus_comarca_born | Centroid_X(LON) | Centroid_Y(LAT) | Propositus_Assigned_comarca | Coded_comarca | Propositus_Assigned_Bishopric | Coded_bishopric | Bishopric_Father | Comarca_Father | Bishopric_Mother | Comarca_Mother | Bishopric_GF__Father | Comarca_GF__Father | Bishopric_GM__Father | Comarca_GM__Father | Bishopric_GF_Mother | Comarca_GF_Mother | Bishopric_GM_Mother | Comarca_GM_Mother | excluded/included |
|---|---|---|---|---|---|---|---|---|---|---|---|---|---|---|---|---|---|---|---|---|---|---|---|---|---|
| H1 | Female | 54 | Berga | Hospital de Campdevànol | Berguedà | Berguedà | 1,807901 | 42,131904 | Berguedà | BBG | Solsona | SOL | Solsona | Berguedà | Solsona | Berguedà | Solsona | Berguedà | Solsona | Berguedà | Solsona | Berguedà | Solsona | Berguedà | included |
| H100 | Male | 18 | Berga | Hospital de Campdevànol | Berguedà | Berguedà | 1,765872 | 42,041683 | Berguedà | BBG | Solsona | SOL | Solsona | Berguedà | Solsona | Berguedà | Solsona | Berguedà | Solsona | Berguedà | Solsona | Berguedà | Solsona | Berguedà | included |
| H101 | Male | 59 | Berga | Hospital de Campdevànol | Berguedà | Berguedà | 1,844025 | 42,103289 | Berguedà | BBG | Solsona | SOL | Solsona | Berguedà | Solsona | Berguedà | Solsona | Berguedà | Solsona | Berguedà | Solsona | Berguedà | Solsona | Berguedà | included |
| H102 | Female | 62 | Berga | Hospital de Campdevànol | Berguedà | Berguedà | 1,932979 | 42,082087 | Berguedà | BBG | Solsona | SOL | Solsona | Berguedà | Solsona | Berguedà | Solsona | Berguedà | Solsona | Berguedà | Solsona | Berguedà | Solsona | Berguedà | included |
| H103 | Male | 24 | Berga | Hospital de Campdevànol | Berguedà | Berguedà | 1,864297 | 42,162698 | Berguedà | BBG | Solsona | SOL | Solsona | Berguedà | Solsona | Berguedà | Solsona | Berguedà | Solsona | Berguedà | Solsona | Berguedà | Solsona | Berguedà | included |
| H104 | Female | 56 | Berga | Hospital de Campdevànol | Berguedà | Berguedà | 1,882196 | 42,103553 | Berguedà | BBG | Solsona | SOL | Solsona | Berguedà | Solsona | Berguedà | Solsona | Berguedà | Solsona | Ripollès | Solsona | Berguedà | Solsona | Berguedà | included |
| H106 | Male | 54 | Berga | Hospital de Campdevànol | Berguedà | Berguedà | 1,825682 | 42,10585 | Berguedà | BBG | Solsona | SOL | Solsona | Berguedà | Solsona | Berguedà | Solsona | Berguedà | Solsona | Berguedà | Solsona | Berguedà | Solsona | Berguedà | included |
| H107 | Female | 65 | Berga | Hospital de Campdevànol | Berguedà | Berguedà | 1,997691 | 42,14624 | Berguedà | BBG | Solsona | SOL | Solsona | Berguedà | Solsona | Berguedà | Solsona | Berguedà | Solsona | Ripollès | Solsona | Berguedà | Solsona | Ripollès | included |
| H111 | Female | 66 | Berga | Hospital de Campdevànol | Berguedà | Berguedà | 1,901261 | 42,047699 | Berguedà | BBG | Solsona | SOL | Solsona | Berguedà | Solsona | Berguedà | Solsona | Berguedà | Solsona | Berguedà | Solsona | Berguedà | Solsona | Berguedà | included |
| H114 | Male | 70 | Berga | Hospital de Campdevànol | Berguedà | Berguedà | 1,776498 | 42,097422 | Berguedà | BBG | Solsona | SOL | Solsona | Berguedà | Solsona | Berguedà | Solsona | Berguedà | Solsona | Berguedà | Solsona | Berguedà | Solsona | Berguedà | excluded (kinship ≥0.09 ) |
| H121 | Female | 54 | Berga | Hospital de Campdevànol | Berguedà | Berguedà | 1,758664 | 42,008776 | Berguedà | BBG | Solsona | SOL | Solsona | Berguedà | Solsona | Berguedà | Solsona | Berguedà | Solsona | Berguedà | Solsona | Berguedà | Vic | Anoia | included |
| H123 | Male | NA | Berga | Hospital de Campdevànol | Berguedà | Berguedà | 1,535513 | 42,126375 | Solsonès | BBG | Solsona | SOL | Urgell | Alt Urgell | Solsona | Berguedà | Urgell | Alt Urgell | Urgell | Alt Urgell | Solsona | Berguedà | Solsona | Berguedà | included |
| H126 | Female | 56 | Berga | Hospital de Campdevànol | Berguedà | Berguedà | 1,833673 | 42,107942 | Berguedà | BBG | Solsona | SOL | Solsona | Berguedà | Solsona | Berguedà | Solsona | Berguedà | Solsona | Berguedà | Solsona | Berguedà | Solsona | Berguedà | included |
| H128 | Female | 58 | Berga | Hospital de Campdevànol | Berguedà | Berguedà | 1,781839 | 42,24195 | Berguedà | BBG | Solsona | SOL | Solsona | Berguedà | Solsona | Berguedà | Solsona | Berguedà | Solsona | Berguedà | Solsona | Berguedà | Solsona | Berguedà | included |
| H129 | Male | 36 | Berga | Hospital de Campdevànol | Berguedà | Berguedà | 1,871052 | 41,998817 | Berguedà | BBG | Solsona | SOL | Solsona | Berguedà | Solsona | Berguedà | Solsona | Berguedà | Solsona | Berguedà | Solsona | Berguedà | Solsona | Berguedà | included |
| H13 | Female | 60 | Berga | Hospital de Campdevànol | Berguedà | Berguedà | 1,825682 | 42,10585 | Berguedà | BBG | Solsona | SOL | Solsona | Berguedà | Vic | Osona | Solsona | Berguedà | Solsona | Berguedà | Solsona | Berguedà | Solsona | Berguedà | included |
| H130 | Male | 55 | Berga | Hospital de Campdevànol | Berguedà | Berguedà | 1,825682 | 42,10585 | Berguedà | BBG | Solsona | SOL | Solsona | Berguedà | Solsona | Berguedà | Solsona | Berguedà | Solsona | Berguedà | Solsona | Berguedà | Solsona | Berguedà | included |
| H132 | Female | 28 | Berga | Hospital de Campdevànol | Berguedà | Berguedà | 1,722596 | 42,065661 | Berguedà | BBG | Solsona | SOL | Solsona | Berguedà | Solsona | Solsonès | Solsona | Berguedà | Solsona | Berguedà | Solsona | Solsonès | Solsona | Berguedà | included |
| H133 | Female | 60 | Berga | Hospital de Campdevànol | Berguedà | Berguedà | 1,857471 | 42,012348 | Berguedà | BBG | Solsona | SOL | Solsona | Berguedà | Solsona | Berguedà | Solsona | Berguedà | Solsona | Berguedà | Solsona | Berguedà | Solsona | Berguedà | included |
| H134 | Male | 68 | Berga | Hospital de Campdevànol | Berguedà | Berguedà | 1,928727 | 42,137344 | Berguedà | BBG | Solsona | SOL | Solsona | Berguedà | Solsona | Berguedà | Solsona | Berguedà | Solsona | Berguedà | Solsona | Berguedà | Solsona | Berguedà | included |
| H135 | Female | 59 | Berga | Hospital de Campdevànol | Berguedà | Berguedà | 1,844025 | 42,103289 | Berguedà | BBG | Solsona | SOL | Solsona | Berguedà | Solsona | Berguedà | Solsona | Berguedà | Solsona | Berguedà | Solsona | Berguedà | Solsona | Berguedà | included |
| H137 | Male | 59 | Berga | Hospital de Campdevànol | Berguedà | Berguedà | 1,834666 | 42,090349 | Berguedà | BBG | Solsona | SOL | Solsona | Berguedà | Solsona | Berguedà | Solsona | Berguedà | Solsona | Berguedà | Solsona | Berguedà | Solsona | Berguedà | included |
| H139 | Male | 41 | Berga | Hospital de Campdevànol | Berguedà | Berguedà | 1,749068 | 42,013148 | Berguedà | BBG | Solsona | SOL | Solsona | Berguedà | Solsona | Bages | Solsona | Berguedà | Solsona | Berguedà | Solsona | Bages | Solsona | Berguedà | included |
| [illegible] | | | | | | | | | | | | | | | | | | | | | | | | | |

| Comarca | Coded_comarca |
|---|---|
| Alt Urgell, Noguera, Cerdanya, Andorrra | AUC |
| Berguedà, Solsonès | BBG |
| Garrotxa | GAR |
| Pallars Jussà, Pallars Sobirà, Alta Ribagorça, La Ribagorza | PAR |
| Ripollès, Osona | RIP |

| Bishopric | Coded_bishopric |
|---|---|
| Girona | GIR |
| Lleida | LLE |
| Solsona | SOL |
| Urgell | URG |
| Vic | VIC |

| DNA_code | gender | age_at_recruitment | Recruitment_place | Recruitment_center | Recruitment_comarca | Propositus_comarca_born | Centroid_X(LON) | Centroid_Y(LAT) | Propositus_Assigned_comarca | Coded_comarca | Propositus_Assigned_Bishopric | Coded_bishopric | Bishopric_Father | Comarca_Father | Bishopric_Mother | Comarca_Mother | Bishopric_GF__Father | Comarca_GF__Father | Bishopric_GM__Father | Comarca_GM__Father | Bishopric_GF_Mother | Comarca_GF_Mother | Bishopric_GM_Mother | Comarca_GM_Mother | excluded/included |
|---|---|---|---|---|---|---|---|---|---|---|---|---|---|---|---|---|---|---|---|---|---|---|---|---|---|
| S26 | Male | 52 | La Seu d'Urgell | Fundació Sant Hospital La Seu d’Urgell | Alt Urgell | Alt Urgell | 1,352228 | 42,366094 | Alt Urgell | AUC | Urgell | URG | Urgell | Alt Urgell | Urgell | Alt Urgell | Urgell | Alt Urgell | Urgell | Alt Urgell | Urgell | Alt Urgell | Urgell | Alt Urgell | included |
| S27 | Female | 43 | La Seu d'Urgell | Fundació Sant Hospital La Seu d’Urgell | Alt Urgell | Alt Urgell | 1,358567 | 42,269368 | Alt Urgell | AUC | Urgell | URG | Urgell | Alt Urgell | Urgell | Alt Urgell | Urgell | Alt Urgell | Urgell | Alt Urgell | Urgell | Alt Urgell | Urgell | Alt Urgell | included |
| S28 | Male | 52 | La Seu d'Urgell | Fundació Sant Hospital La Seu d’Urgell | Alt Urgell | Alt Urgell | 1,455904 | 42,359941 | Alt Urgell | AUC | Urgell | URG | Urgell | Alt Urgell | Urgell | Alt Urgell | Urgell | Alt Urgell | Urgell | Alt Urgell | Urgell | Alt Urgell | Urgell | Alt Urgell | included |
| S29 | Male | 57 | La Seu d'Urgell | Fundació Sant Hospital La Seu d’Urgell | Alt Urgell | Alt Urgell | 1,469832 | 42,21881 | Alt Urgell | AUC | Urgell | URG | Urgell | Alt Urgell | Urgell | Alt Urgell | Urgell | Alt Urgell | Urgell | Alt Urgell | Urgell | Alt Urgell | Urgell | Alt Urgell | included |
| S3 | Female | 27 | La Seu d'Urgell | Fundació Sant Hospital La Seu d’Urgell | Alt Urgell | Alt Urgell | 1,743658 | 42,385349 | Cerdanya | AUC | Urgell | URG | Urgell | Cerdanya | Urgell | Cerdanya | Urgell | Cerdanya | Urgell | Cerdanya | Urgell | Cerdanya | Urgell | Cerdanya | included |
| S30 | Female | 63 | La Seu d'Urgell | Fundació Sant Hospital La Seu d’Urgell | Alt Urgell | Alt Urgell | 1,483087 | 42,337753 | Alt Urgell | AUC | Urgell | URG | Urgell | Alt Urgell | Urgell | Alt Urgell | Urgell | Alt Urgell | Urgell | Cerdanya | Urgell | Alt Urgell | Urgell | Alt Urgell | included |
| S32 | Female | 69 | La Seu d'Urgell | Fundació Sant Hospital La Seu d’Urgell | Alt Urgell | Alt Urgell | 1,325459 | 42,461829 | Pallars Sobirà | PAR | Urgell | URG | Urgell | Alt Urgell | Urgell | Pallars Sobirà | Urgell | Alt Urgell | Urgell | Alt Urgell | Urgell | Pallars Sobirà | Urgell | Pallars Sobirà | included |
| S33 | Female | 44 | La Seu d'Urgell | Fundació Sant Hospital La Seu d’Urgell | Alt Urgell | Alt Urgell | 1,497603 | 42,338351 | Alt Urgell | AUC | Urgell | URG | Urgell | Cerdanya | Urgell | Cerdanya | Urgell | Cerdanya | Urgell | Alt Urgell | Urgell | Alt Urgell | Urgell | Alt Urgell | included |
| S34 | Female | 51 | La Seu d'Urgell | Fundació Sant Hospital La Seu d’Urgell | Alt Urgell | Alt Urgell | 1,316892 | 42,450894 | Pallars Sobirà | PAR | Urgell | URG | Urgell | Alt Urgell | Urgell | Pallars Sobirà | ND | ND | Urgell | Alt Urgell | Urgell | Pallars Sobirà | Urgell | Pallars Sobirà | included |
| S35 | Male | 54 | La Seu d'Urgell | Fundació Sant Hospital La Seu d’Urgell | Alt Urgell | Pallars Jussà | 0,864251 | 42,450711 | Pallars Jussà | PAR | Lleida | LLE | Urgell | Alta Ribagorça | Urgell | Pallars Jussà | Urgell | Alta Ribagorça | Urgell | Alta Ribagorça | Lleida | Alta Ribagorça | Urgell | Pallars Jussà | included |
| S37 | Female | 41 | La Seu d'Urgell | Fundació Sant Hospital La Seu d’Urgell | Alt Urgell | Alt Urgell | 1,313166 | 42,324396 | Alt Urgell | AUC | Urgell | URG | Urgell | Alt Urgell | Urgell | Alt Urgell | Urgell | Pallars Sobirà | Urgell | Pallars Sobirà | Urgell | Alt Urgell | Urgell | Alt Urgell | included |
| S38 | Female | 39 | La Seu d'Urgell | Fundació Sant Hospital La Seu d’Urgell | Alt Urgell | Alt Urgell | 1,287186 | 42,175174 | Alt Urgell | AUC | Urgell | URG | Urgell | Alt Urgell | Vic | Bages | Urgell | Alt Urgell | Urgell | Alt Urgell | Barcelona | Barcelonès | Urgell | Alt Urgell | included |
| S39 | Female | 52 | La Seu d'Urgell | Fundació Sant Hospital La Seu d’Urgell | Alt Urgell | Alt Urgell | 1,471884 | 42,268966 | Alt Urgell | AUC | Urgell | URG | Urgell | Alt Urgell | Urgell | Alt Urgell | Solsona | Solsonès | Urgell | Alt Urgell | Urgell | Alt Urgell | Urgell | Alt Urgell | included |
| S4 | Female | 47 | La Seu d'Urgell | Fundació Sant Hospital La Seu d’Urgell | Alt Urgell | Alt Urgell | 1,609127 | 42,35824 | Alt Urgell | AUC | Urgell | URG | Urgell | Alt Urgell | Urgell | Cerdanya | Urgell | Alt Urgell | Urgell | Alt Urgell | Urgell | Cerdanya | Urgell | Cerdanya | included |
| S40 | Male | 70 | La Seu d'Urgell | Fundació Sant Hospital La Seu d’Urgell | Alt Urgell | Cerdanya | 1,843059 | 42,41224 | Cerdanya | AUC | Urgell | URG | Urgell | Cerdanya | Urgell | Cerdanya | Urgell | Cerdanya | Urgell | Cerdanya | Urgell | Cerdanya | Urgell | Cerdanya | excluded (kinship ≥0.09 ) |
| S41 | Male | 38 | La Seu d'Urgell | Fundació Sant Hospital La Seu d’Urgell | Alt Urgell | Alt Urgell | 1,285417 | 42,283932 | Alt Urgell | AUC | Urgell | URG | Urgell | Alt Urgell | Urgell | Alt Urgell | Urgell | Alt Urgell | Urgell | Alt Urgell | Urgell | Alt Urgell | Urgell | Pallars Jussà | excluded (kinship ≥0.09 ) |
| S42 | Male | 43 | La Seu d'Urgell | Fundació Sant Hospital La Seu d’Urgell | Alt Urgell | Pallars Jussà | 1,065978 | 42,383139 | Pallars Sobirà | PAR | Urgell | URG | Urgell | Pallars Sobirà | Urgell | Pallars Jussà | Urgell | Pallars Sobirà | Urgell | Pallars Sobirà | Urgell | Pallars Jussà | Urgell | Pallars Jussà | included |
| S43 | Female | 48 | La Seu d'Urgell | Fundació Sant Hospital La Seu d’Urgell | Alt Urgell | Alt Urgell | 1,319347 | 42,376299 | Alt Urgell | AUC | Urgell | URG | Urgell | Alt Urgell | Urgell | Alt Urgell | Urgell | Alt Urgell | Urgell | Alt Urgell | Urgell | Alt Urgell | Urgell | Pallars Sobirà | included |
| S44 | Female | 62 | La Seu d'Urgell | Fundació Sant Hospital La Seu d’Urgell | Alt Urgell | Pallars Jussà | 1,091762 | 42,374859 | Pallars Sobirà | PAR | Urgell | URG | Urgell | Pallars Jussà | Urgell | Pallars Sobirà | Urgell | Pallars Jussà | Urgell | Pallars Jussà | Urgell | Pallars Sobirà | Urgell | Pallars Sobirà | included |
| S45 | Female | 51 | La Seu d'Urgell | Fundació Sant Hospital La Seu d’Urgell | Alt Urgell | Alt Urgell | 1,42027 | 42,3169 | Alt Urgell | AUC | Urgell | URG | Urgell | Alt Urgell | Urgell | Alt Urgell | Urgell | Alt Urgell | Urgell | Alt Urgell | Urgell | Alt Urgell | Urgell | Alt Urgell | included |
| S46 | Female | 38 | La Seu d'Urgell | Fundació Sant Hospital La Seu d’Urgell | Alt Urgell | Alt Urgell | 1,43067 | 42,363381 | Alt Urgell | AUC | Urgell | URG | Urgell | Alt Urgell | Urgell | Alt Urgell | Urgell | Alt Urgell | Urgell | Alt Urgell | Urgell | Alt Urgell | Urgell | Alt Urgell | excluded (kinship ≥0.09 ) |
| S48 | Female | 49 | La Seu d'Urgell | Fundació Sant Hospital La Seu d’Urgell | Alt Urgell | Alt Urgell | 1,461979 | 42,357709 | Alt Urgell | AUC | Urgell | URG | Urgell | Alt Urgell | Urgell | Alt Urgell | Urgell | Alt Urgell | Urgell | Alt Urgell | Urgell | Alt Urgell | Urgell | Alt Urgell | included |
| S49 | Male | 57 | La Seu d'Urgell | Fundació Sant Hospital La Seu d’Urgell | Alt Urgell | Alt Urgell | 1,648503 | 42,362204 | Alt Urgell | AUC | Urgell | URG | Urgell | Alt Urgell | Urgell | Cerdanya | Urgell | Alt Urgell | Urgell | Alt Urgell | Urgell | Cerdanya | Urgell | Cerdanya | included |
| S50 | Female | 64 | La Seu d'Urgell | Fundació Sant Hospital La Seu d’Urgell | Alt Urgell | Alt Urgell | 1,483087 | 42,337753 | Alt Urgell | AUC | Urgell | URG | Urgell | Alt Urgell | Urgell | Alt Urgell | Urgell | Alt Urgell | Urgell | Cerdanya | Urgell | Alt Urgell | Urgell | Alt Urgell | excluded (kinship ≥0.09 ) |
| S52 | Male | 52 | La Seu d'Urgell | Fundació Sant Hospital La Seu d’Urgell | Alt Urgell | ND | 1,531975 | 42,367062 | Alt Urgell | AUC | Urgell | URG | Urgell | Alt Urgell | Urgell | Alt Urgell | Urgell | Alt Urgell | Urgell | Alt Urgell | Urgell | Alt Urgell | Urgell | Alt Urgell | included |
| S53 | Male | 60 | La Seu d'Urgell | Fundació Sant Hospital La Seu d’Urgell | Alt Urgell | Pallars Jussà | 0,806515 | 42,366431 | Alta Ribagorça | PAR | Lleida | LLE | Lleida | Alta Ribagorça | Urgell | Pallars Jussà | Lleida | Alta Ribagorça | Lleida | Alta Ribagorça | Lleida | Alta Ribagorça | Urgell | Pallars Jussà | excluded (kinship ≥0.09 ) |
| S54 | Female | 57 | La Seu d'Urgell | Fundació Sant Hospital La Seu d’Urgell | Alt Urgell | Alt Urgell | 1,283313 | 42,059888 | Alt Urgell | AUC | Urgell | URG | Urgell | Alt Urgell | Urgell | Alt Urgell | ND | ND | Urgell | Alt Urgell | Urgell | Alt Urgell | Urgell | Alt Urgell | included |
| S56 | Male | 63 | La Seu d'Urgell | Fundació Sant Hospital La Seu d’Urgell | Alt Urgell | Pallars Jussà | 1,331988 | 42,31041 | Alt Urgell | AUC | Urgell | URG | Urgell | Alt Urgell | Urgell | Pallars Jussà | Urgell | Alt Urgell | Urgell | Alt Urgell | Urgell | Pallars Jussà | Urgell | Pallars Jussà | excluded (kinship ≥0.09 ) |
| S57 | Female | 55 | La Seu d'Urgell | Fundació Sant Hospital La Seu d’Urgell | Alt Urgell | ND | 1,408592 | 42,444172 | Alt Urgell | AUC | Urgell | URG | Urgell | Alt Urgell | Urgell | Alt Urgell | Urgell | Alt Urgell | Urgell | Alt Urgell | Urgell | Alt Urgell | Urgell | Alt Urgell | excluded (kinship ≥0.09 ) |
| S58 | Male | 63 | La Seu d'Urgell | Fundació Sant Hospital La Seu d’Urgell | Alt Urgell | Pallars Sobirà | 1,114305 | 42,486797 | Pallars Sobirà | PAR | Urgell | URG | Urgell | Pallars Sobirà | Urgell | Pallars Sobirà | Urgell | Pallars Sobirà | Urgell | Pallars Sobirà | Urgell | Pallars Sobirà | Urgell | Pallars Sobirà | included |
| S59 | Male | 34 | La Seu d'Urgell | Fundació Sant Hospital La Seu d’Urgell | Alt Urgell | Alt Urgell | 1,3273 | 42,280308 | Alt Urgell | AUC | Urgell | URG | Urgell | Alt Urgell | Urgell | Alt Urgell | Urgell | Alt Urgell | Urgell | Alt Urgell | Urgell | Alt Urgell | Urgell | Alt Urgell | excluded (kinship ≥0.09 ) |
| S61 | Female | 37 | La Seu d'Urgell | Fundació Sant Hospital La Seu d’Urgell | Alt Urgell | Alt Urgell | 1,743658 | 42,385349 | Cerdanya | AUC | Urgell | URG | Urgell | Cerdanya | Urgell | Cerdanya | Urgell | Cerdanya | Urgell | Cerdanya | Urgell | Cerdanya | Urgell | Cerdanya | excluded (kinship ≥0.09 ) |
| S62 | Female | 64 | La Seu d'Urgell | Fundació Sant Hospital La Seu d’Urgell | Alt Urgell | Alt Urgell | 1,528045 | 42,306146 | Alt Urgell | AUC | Urgell | URG | Urgell | Alt Urgell | Urgell | Alt Urgell | Urgell | Cerdanya | Urgell | Alt Urgell | Urgell | Alt Urgell | Urgell | Alt Urgell | excluded (kinship ≥0.09 ) |
| S64 | Male | 49 | La Seu d'Urgell | Fundació Sant Hospital La Seu d’Urgell | Alt Urgell | Alt Urgell | 1,451773 | 42,375788 | Alt Urgell | AUC | Urgell | URG | Urgell | Alt Urgell | Urgell | Alt Urgell | Urgell | Alt Urgell | Urgell | Alt Urgell | Urgell | Alt Urgell | Urgell | Alt Urgell | included |
| S65 | Male | 70 | La Seu d'Urgell | Fundació Sant Hospital La Seu d’Urgell | Alt Urgell | Alt Urgell | 1,195361 | 42,509101 | Pallars Sobirà | PAR | Urgell | URG | Urgell | Pallars Sobirà | Urgell | Alt Urgell | Urgell | Pallars Sobirà | Urgell | Pallars Sobirà | Urgell | Alt Urgell | Urgell | Alt Urgell | included |
| S66 | Female | 50 | La Seu d'Urgell | Fundació Sant Hospital La Seu d’Urgell | Alt Urgell | Alt Urgell | 1,510139 | 42,174769 | Solsonès | BBG | Urgell | URG | Urgell | Alt Urgell | Urgell | Alt Urgell | Urgell | Alt Urgell | Vic | Bages | Urgell | Alt Urgell | Urgell | Alt Urgell | included |
| S68 | Female | 56 | La Seu d'Urgell | Fundació Sant Hospital La Seu d’Urgell | Alt Urgell | Alt Urgell | 1,056629 | 42,28328 | Pallars Sobirà | PAR | Urgell | URG | Urgell | Alt Urgell | Urgell | Pallars Jussà | Urgell | Pallars Jussà | Urgell | Alt Urgell | Urgell | Pallars Jussà | Urgell | Pallars Sobirà | included |
| S69 | Male | 65 | La Seu d'Urgell | Fundació Sant Hospital La Seu d’Urgell | Alt Urgell | Alt Urgell | 1,220872 | 42,476089 | Pallars Sobirà | PAR | Urgell | URG | Urgell | Pallars Sobirà | Urgell | Alt Urgell | Urgell | Pallars Sobirà | Urgell | Pallars Sobirà | Urgell | Alt Urgell | Urgell | Alt Urgell | excluded (kinship ≥0.09 ) |
| S73 | Female | 38 | La Seu d'Urgell | Fundació Sant Hospital La Seu d’Urgell | Alt Urgell | Alt Urgell | 1,348944 | 42,177788 | Alt Urgell | AUC | Urgell | URG | Urgell | Alt Urgell | Urgell | Alt Urgell | Urgell | Alt Urgell | Urgell | Alt Urgell | Urgell | Alt Urgell | Urgell | Alt Urgell | included |
| S75 | Male | 69 | La Seu d'Urgell | Fundació Sant Hospital La Seu d’Urgell | Alt Urgell | Cerdanya | 1,849125 | 42,401554 | Cerdanya | AUC | Urgell | URG | Urgell | Cerdanya | Urgell | Cerdanya | Urgell | Cerdanya | Urgell | Cerdanya | Urgell | Cerdanya | Urgell | Cerdanya | included |
| S76 | Female | 48 | La Seu d'Urgell | Fundació Sant Hospital La Seu d’Urgell | Alt Urgell | Alt Urgell | 1,52991 | 42,367823 | Alt Urgell | AUC | Urgell | URG | Urgell | Alt Urgell | Urgell | Alt Urgell | Urgell | Alt Urgell | Urgell | Alt Urgell | Urgell | Alt Urgell | Urgell | Alt Urgell | included |
| S77 | Male | 50 | La Seu d'Urgell | Fundació Sant Hospital La Seu d’Urgell | Alt Urgell | Alt Urgell | 1,476875 | 42,411588 | Alt Urgell | AUC | Urgell | URG | Urgell | Alt Urgell | Urgell | Alt Urgell | Urgell | Alt Urgell | Urgell | Alt Urgell | Urgell | Alt Urgell | Urgell | Andorra | included |
| S78 | Female | 37 | La Seu d'Urgell | Fundació Sant Hospital La Seu d’Urgell | Alt Urgell | Alt Urgell | 1,346046 | 42,385458 | Alt Urgell | AUC | Urgell | URG | Urgell | Alt Urgell | Urgell | Alt Urgell | Urgell | Alt Urgell | Urgell | Alt Urgell | Urgell | Alt Urgell | Urgell | Pallars Sobirà | excluded (kinship ≥0.09 ) |
| S79 | Male | 66 | La Seu d'Urgell | Fundació Sant Hospital La Seu d’Urgell | Alt Urgell | Alt Urgell | 1,760651 | 42,217094 | Berguedà | BBG | Solsona | SOL | Urgell | Alt Urgell | Vic | Berguedà | Urgell | Alt Urgell | Urgell | Alt Urgell | Urgell | Pallars Jussà | Urgell | Pallars Jussà | included |
| S8 | Female | 56 | La Seu d'Urgell | Fundació Sant Hospital La Seu d’Urgell | Alt Urgell | Alt Urgell | 1,583392 | 42,264886 | Alt Urgell | AUC | Urgell | URG | Urgell | Alt Urgell | Urgell | Alt Urgell | Urgell | Alt Urgell | Urgell | Alt Urgell | Urgell | Alt Urgell | Urgell | Alt Urgell | included |
| S81 | Female | 39 | La Seu d'Urgell | Fundació Sant Hospital La Seu d’Urgell | Alt Urgell | Alt Urgell | 1,305924 | 42,234796 | Alt Urgell | AUC | Urgell | URG | Urgell | Alt Urgell | Urgell | Alt Urgell | Urgell | Alt Urgell | Urgell | Pallars Jussà | Urgell | Alt Urgell | Urgell | Alt Urgell | excluded (kinship ≥0.09 ) |
| S82 | Male | 37 | La Seu d'Urgell | Fundació Sant Hospital La Seu d’Urgell | Alt Urgell | Alt Urgell | 1,50706 | 42,337605 | Alt Urgell | AUC | Urgell | URG | Urgell | Alt Urgell | Barcelona | Barcelonès | Urgell | Alt Urgell | Urgell | Alt Urgell | Barcelona | Barcelonès | Urgell | Alt Urgell | excluded (kinship ≥0.09 ) |
| S83 | Female | 39 | La Seu d'Urgell | Fundació Sant Hospital La Seu d’Urgell | Alt Urgell | Alt Urgell | 1,409579 | 42,382484 | Alt Urgell | AUC | Urgell | URG | Urgell | Alt Urgell | Urgell | Alt Urgell | Urgell | Alt Urgell | Urgell | Alt Urgell | Urgell | Alt Urgell | Barcelona | Barcelonès | included |
| S84 | Female | 29 | La Seu d'Urgell | Fundació Sant Hospital La Seu d’Urgell | Alt Urgell | Alt Urgell | 1,448227 | 42,344318 | Alt Urgell | AUC | Urgell | URG | Urgell | Alt Urgell | Urgell | Alt Urgell | Urgell | Alt Urgell | Urgell | Pallars Jussà | Urgell | Alt Urgell | Urgell | Alt Urgell | included |
| S85 | Male | 53 | La Seu d'Urgell | Fundació Sant Hospital La Seu d’Urgell | Alt Urgell | Alt Urgell | 1,348308 | 42,377453 | Alt Urgell | AUC | Urgell | URG | Urgell | Alt Urgell | Urgell | Alt Urgell | Urgell | Alt Urgell | Urgell | Pallars Sobirà | Urgell | Alt Urgell | Urgell | Alt Urgell | included |
| S86 | Female | 51 | La Seu d'Urgell | Fundació Sant Hospital La Seu d’Urgell | Alt Urgell | Alt Urgell | 1,373782 | 42,320818 | Alt Urgell | AUC | Urgell | URG | Urgell | Alt Urgell | Urgell | Alt Urgell | Urgell | Alt Urgell | Urgell | Alt Urgell | Urgell | Alt Urgell | Urgell | Pallars Sobirà | included |
| S87 | Male | 62 | La Seu d'Urgell | Fundació Sant Hospital La Seu d’Urgell | Alt Urgell | Alt Urgell | 2,060527 | 42,190346 | Berguedà | BBG | Solsona | SOL | Urgell | Alt Urgell | Girona | Garrotxa | Vic | Ripollès | Urgell | Ripollès | Girona | Garrotxa | Urgell | Alt Urgell | excluded (kinship ≥0.09 ) |
| S88 | Female | 44 | La Seu d'Urgell | Fundació Sant Hospital La Seu d’Urgell | Alt Urgell | Alt Urgell | 1,478377 | 42,382915 | Alt Urgell | AUC | Urgell | URG | Urgell | Alt Urgell | Urgell | Alt Urgell | Urgell | Alt Urgell | Urgell | Alt Urgell | Urgell | Alt Urgell | Urgell | Alt Urgell | included |
| S89 | Female | 59 | La Seu d'Urgell | Fundació Sant Hospital La Seu d’Urgell | Alt Urgell | Alt Urgell | 1,233729 | 42,376579 | Pallars Sobirà | PAR | Urgell | URG | Urgell | Alt Urgell | Urgell | Pallars Sobirà | Urgell | Alt Urgell | Urgell | Pallars Sobirà | Urgell | Pallars Sobirà | Urgell | Alt Urgell | included |
| S9 | Female | NA | La Seu d'Urgell | Fundació Sant Hospital La Seu d’Urgell | Alt Urgell | Ripollès | 2,166507 | 42,316868 | Ripollès | RIP | Urgell | URG | Urgell | Ripollès | Urgell | Ripollès | Urgell | Ripollès | Urgell | Ripollès | Urgell | Ripollès | Urgell | Ripollès | excluded (kinship ≥0.09 ) |
| S90 | Female | 21 | La Seu d'Urgell | Fundació Sant Hospital La Seu d’Urgell | Alt Urgell | Alt Urgell | 1,413557 | 42,310427 | Alt Urgell | AUC | Urgell | URG | Urgell | Alt Urgell | Urgell | Alt Urgell | Urgell | Alt Urgell | Barcelona | Vallés Occidental | Urgell | Alt Urgell | Urgell | Alt Urgell | included |
| S93 | Female | 64 | La Seu d'Urgell | Fundació Sant Hospital La Seu d’Urgell | Alt Urgell | Alt Urgell | 1,348917 | 42,480647 | Alt Urgell | AUC | Urgell | URG | Urgell | Pallars Sobirà | Urgell | Alt Urgell | Urgell | Pallars Sobirà | Urgell | Pallars Sobirà | Urgell | Alt Urgell | Urgell | Alt Urgell | included |
| S94 | Male | 54 | La Seu d'Urgell | Fundació Sant Hospital La Seu d’Urgell | Alt Urgell | Alt Urgell | 1,69742 | 42,35105 | Cerdanya | AUC | Urgell | URG | Urgell | Cerdanya | Urgell | Cerdanya | Urgell | Cerdanya | Urgell | Cerdanya | Urgell | Alt Urgell | Urgell | Cerdanya | included |
| S96 | Male | 48 | La Seu d'Urgell | Fundació Sant Hospital La Seu d’Urgell | Alt Urgell | Alt Urgell | 1,363726 | 42,284145 | Alt Urgell | AUC | Urgell | URG | Urgell | Alt Urgell | Urgell | Alt Urgell | Urgell | Alt Urgell | Urgell | Alt Urgell | Urgell | Alt Urgell | Urgell | Alt Urgell | included |
| S99 | Female | 38 | La Seu d'Urgell | Fundació Sant Hospital La Seu d’Urgell | Alt Urgell | Alt Urgell | 1,358567 | 42,269368 | Alt Urgell | AUC | Urgell | URG | Urgell | Alt Urgell | Urgell | Alt Urgell | Urgell | Alt Urgell | Urgell | Alt Urgell | Urgell | Alt Urgell | Urgell | Alt Urgell | excluded (kinship ≥0.09 ) |

| Comarca | Coded_comarca |
|---|---|
